# The mycobacterial guaB1 gene encodes a guanosine 5’-monophosphate reductase with a cystathione-β-synthase domain

**DOI:** 10.1101/2021.08.05.455185

**Authors:** Zdeněk Knejzlík, Michal Doležal, Klára Herkommerová, Kamila Clarova, Martin Klíma, Matteo Dedola, Eva Zborníková, Dominik Rejman, Iva Pichová

## Abstract

Purine metabolism plays a pivotal role in bacterial life cycle, however, regulation of the *de novo* and purine salvage pathways have not been extensively detailed in mycobacteria. By gene knockout, biochemical and structural analyses, we identified *Mycobacterium smegmatis* (Msm) and *Mycobacterium tuberculosis* (Mtb) *guaB1* gene product as a novel type of guanosine 5’-monophosphate reductase (GMPR), which recycles guanosine monophosphate to inosine monophosphate within the purine salvage pathway and contains cystathione β-synthase (CBS) domains with atypical orientation in the octamer. CBS domains share a much larger interacting area with a conserved catalytic domain in comparison with the only known CBS containing protozoan GMPR and closely related inosine monophosphate dehydrogenase structures. Our results revealed essential effect of pH on allosteric regulation of Msm GMPR activity and oligomerization with adenine and guanosine nucleotides binding to CBS domains.Bioinformatic analysis indicated the presence of GMPRs containing CBS domains across the entire *Actinobacteria* phylum.

## Introduction

In most organisms, there are two pathways for nucleotide biosynthesis: 1) the *de novo* pathway, in which nucleotides are synthesized from 5’-phospho-α-D-ribose-1’-diphosphate (PRPP) in a series of reactions; and 2) the salvage pathway, in which nucleotides are recycled after the breakdown of nucleic acids or coenzymes from free purine or pyrimidine bases (el Kouni, 2003). Mycobacteria, including *Mycobacterium smegmatis* (Msm) and *Mycobacterium tuberculosis* (Mtb), express enzymes from both the *de novo* and purine salvage pathways. However, the interdependence and regulation of these processes during different stages of the mycobacterial life cycle remain unclear (Malathi & Ramakrishnan, 1966). Inosine monophosphate (IMP) is a hub metabolite for the biosynthesis of guanosine monophosphate (GMP) and adenosine monophosphate (AMP), and ATP and GTP regulate biosynthesis of GMP and AMP, respectively. This cross-regulation by the end products can slow down biosynthesis of these nucleotide monophosphates (Pimkin *et al*, 2009). In mycobacteria, IMP is converted to xanthosine monophosphate (XMP) by an NAD^+^-dependent inosine monophosphate dehydrogenase (IMPDH) encoded by the *guaB2* gene (*MSMEG_1602* in Msm and *Rv3411c* in Mtb). This reaction is an essential rate-limiting step in the *de novo* biosynthesis of guanine nucleotides (Cox *et al*, 2016; Ducati *et al*, 2011; Gollapalli *et al*, 2010; Hedstrom *et al*, 2011; Chacko *et al*, 2018; Chen *et al*, 2010; Juvale *et al*, 2019; Makowska-Grzyska *et al*, 2015a; Sahu *et al*, 2018; Sassetti *et al*, 2001, 2003; Singh *et al*, 2017; Usha *et al*, 2011). In subsequent reactions, ATP-dependent GMP synthetase catalyzes conversion of XMP to GMP, which can either be further converted to GDP and GTP or salvaged back to IMP by a two-step reaction catalyzed by an NADPH-dependent guanosine 5’-monophosphate dehydrogenase (GMPR). During the first step, GMP is deaminated and the covalent intermediate adduct GMPR-XMP* and NH_3_ are formed. In the second step, the intermediate adduct is reduced to IMP by NADPH, and both reaction products (NAD^+^ and IMP) are released from GMPR (Hedstrom, 2012).

IMPDHs and GMPRs belong to the IMPDH/GMPR enzyme family. The members of this family share several common structural features, including a (β/α)_8_ barrel structure of the catalytic domains and core tetrameric or octameric structural organization. They also share modes of ligand binding and formation of a covalent enzyme-XMP* catalytic intermediate (Hedstrom, 2009; Hedstrom, 2012; Hedstrom & Gan, 2006). A structural feature characteristic of IMPDHs is the presence of cystathionine-β-synthase domains (CBSs; CBS dimers are called Bateman domains), which usually regulate the enzyme activity or multimerization upon nucleotide binding (Ereno-Orbea *et al*, 2013; Gan *et al*, 2002; Makowska-Grzyska *et al*., 2015a; Makowska-Grzyska *et al*, 2015b; Nimmesgern *et al*, 1999). Deletion of the IMPDH CBS in *E. coli* causes dysregulation of the adenine and guanine nucleotide pool (Pimkin & Markham, 2008; Pimkin *et al*., 2009). GMPRs typically do not possess CBSs, with the exception of GMPRs from the pathogenic protozoans *Trypanosoma brucei* (Tb GMPR) (Bessho *et al*, 2016), *Trypanosoma congolense* (Tc GMPR) (Sarwono *et al*, 2017) and *Leshmania donovani* (Ld GMPR) (Smith *et al*, 2016). Recently, structural analysis of Tb GMPR showed that ATP induces octamer dissociation, while guanine nucleotides do not influence the enzyme’s oligomeric state (Imamura *et al*, 2020).

While IMPDH activity is strictly required for guanine nucleotide metabolism, the role of salvaging GMPR reductase activity at different bacterial lifecycle stages and under stress conditions is still not well understood. GMPR is not essential for the growth of either Gram-negative bacteria such as *E. coli* (Magasanik & Karibian, 1960; Patton *et al*, 2011) or Gram-positive species such as *B. subtilis* (Endo *et al*, 1983), and its expression can be regulated by the intracellular ratio of adenine and guanine nucleotide pools (Benson & Gots, 1975; Kessler & Gots, 1985).

The presence of an enzyme with GMPR activity in mycobacteria has not yet been identified. The Msm genome contains three IMPDH homologues: *guaB1* (MSMEG_3634), *guaB2* (MSMEG_1602), and *guaB3* (MSMEG_1603). However, little information is available about the enzymes encoded by these genes. In the Mtb genome, only *guaB2* (Rv3411c) has been shown to encode a functional IMPDH enzyme (Griffin *et al*, 2011; Usha *et al*., 2011), which catalyzes a rate-limiting step in the *de novo* biosynthesis of guanine nucleotides from IMP. Mtb IMPDH inhibition results in the depletion of cellular guanine nucleotides (Hedstrom, 2009).

Here, we show that the Msm and Mtb *guaB1* genes encode an active guanosine 5’-monophosphate reductase, which is not essential for bacterial survival but contributes to recycling of purine nucleotides. The activity of Msm GMPR is allosterically regulated by the ATP:GTP ratio only at pH values under 7, suggesting different GMPR activities in growing bacteria and during latent infection when the intracellular pH decreases (Rao *et al*, 2001; Vandal *et al*, 2009; Zhang *et al*, 1999b). X-ray crystallogrpahy revealed that Msm GMPR contains a CBS domain with an atypical position in the octamer. Phylogenetic analysis indicated the presence of *guaB1*-encoded GMPRs with CBSs throughout the *Actinobacteria* phylum.

## Results

### Msm GuaB1 is an NADPH-dependent GMPR involved in the purine salvage pathway

Msm GuaB1, the enzyme encoded by Msm *guaB1* (MSMEG_3634), shares 92% amino acid sequence similarity with Mtb GuaB1, which is encoded by Mtb *guaB1* (Rv1843c). Random saturating transposon mutagenesis of the Mtb genome showed that *gua B1* is not essential for growth of Mtb (DeJesus *et al*, 2017). To investigate the potential role of GuaB1 in mycobacterial purine nucleotide biosynthesis, we used Msm as a model organism and knocked out the *guaB1* gene. As a control, we also knocked out the *guaB2* gene encoding an IMPDH. The *ΔguaB1* Msm strain did not exhibit any growth defects in media supplemented with hypoxanthine, guanine, and adenine (Fig. 1A), indicating that GuaB1 is not essential for Msm growth. As expected, the *ΔguaB2* strain grew only in medium supplemented with guanine and not in media supplemented with hypoxanthine or adenine (Fig. 1A.), confirming the IMPDH activity of Msm GuaB2.

**Fig 1.**
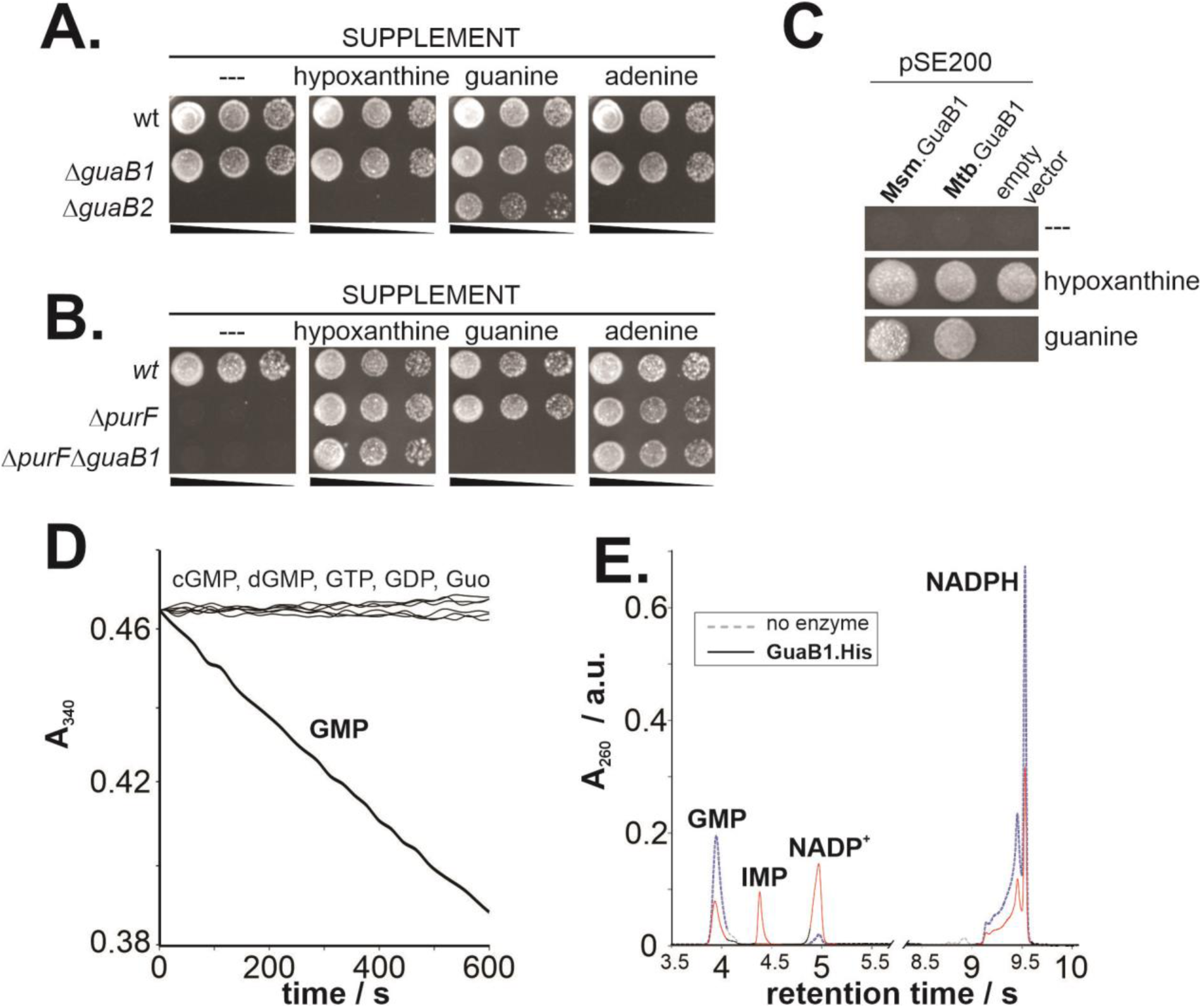
Analysis of Msm GuaB1 function and activity. **A)** Growth of *ΔguaB1* and *ΔguaB2* strains in media supplemented with different purine nucleotides. Exponentially grown Msm strains were spotted at three O.D._600_ values (10^−2^, 10^−3^, and 10^−4^) on 7H10/ADC medium without or with 100 µM purine nucleobase supplement and incubated for 3 days at 37 °C. **B)** Growth of the *ΔpurF and ΔguaB1ΔpurF* Msm strains under the conditions used in panel A. **C)** Complementation of GuaB1 deficiency. The *ΔguaB1ΔpurF* strain transformed with an empty pSE200 plasmid or pSE200 carrying the Msm or Mtb *guaB1* gene was spotted (O.D._600_ = 10^−2^) on 7H10/ADC medium without or with 100 μM purine nucleobase supplement. **D)** Guanosine 5’-monophospate reductase activity of GuaB1 *in vitro.* The absorbance at 340 nm of reaction mixtures containing 200 nM recombinant GuaB1, 75 μM NADPH, and 1 mM guanine nucleotides (cGMP, GMP, dGMP, GDP, GTP) or 1 mM guanosine (Guo) was continuously monitored for 600 s. **F)** Analysis of the reaction mixture with and without GuaB1. Reaction mixtures containing 200 μM NADPH and 100 μM GMP without GuaB1 (dashed blue line) or with 100 nM GuaB1 (full red line) cultivated for 30 min at 25 °C were analyzed by UPLC on a reverse phase column. The peaks were identified based on nucleotide calibration standards.

To assess the effect of *guaB1* gene deletion on purine composition, we analyzed the intracellular purine metabolites in the wt and *ΔguaB1* strains during exponential growth using the fast acetic acid metabolome extraction approach (Varik *et al*, 2017) in combination with hydrophilic interaction liquid chromatography (HILIC) coupled with UV/MS detection (Zbornikova *et al*, 2019). Both strains contained comparable adenine and guanine nucleotide content (Fig. S1). Next, we blocked the *de novo* purine metabolic pathway by knocking out the essential *purF* gene and compared the growth of the *ΔguaB1ΔpurF* strain with that of the wt and *ΔpurF* Msm strains (Knejzlik *et al*, 2020; Knejzlik *et al*, 2019). The *ΔpurF* strain requires for growth an external source of purine such as hypoxanthine, adenine or, guanine to serve as a precursor for purine metabolism *via* the corresponding salvaging enzymes (Knejzlik *et al*., 2020). The *ΔguaB1ΔpurF* strain grew on media containing adenine or hypoxanthine but not on media containing guanine as the sole purine source (Fig. 1B). The defect of guanine salvaging in the *ΔguaB1ΔpurF* strain confirmed the involvement of Msm GuaB1 in nucleotide interconversion to other purines through the purine salvage pathway. Growth of the *ΔguaB1ΔpurF* Msm strain on media supplemented with guanine was restored by complementation with a plasmid encoding Msm or Mtb GuaB1 (Fig. 1C), which demonstrates that both Msm and Mtb GuaB1 function within the guanine salvaging pathway.

To further biochemically characterize GuaB1, we produced recombinant Msm GuaB1 in *E. coli* and the less soluble Mtb GuaB1 in the *ΔguaB1* Msm strain. We tested Msm GuaB1 activity in the presence of NADPH and different guanine nucleotides using a spectrophotometric assay (Fig. 1D). Importantly, we detected activity in the presence of GMP but not in the presence of the other guanine nucleotides tested. We found a similar activity pattern for Mtb GuaB1, which explains why a previous biochemical characterization of Mtb GuaB1 (Usha *et al*., 2011) did not confirm the IMPDH activity of this enzyme. Chromatographic analysis of the enzymatic mixture showed formation of IMP and NAD^+^ (Fig. 1E). Our results confirmed that GuaB1 (hereafter referred to as GMPR) functions as a mycobacterial NADPH-dependent guanosine 5’-monophosphate dehydrogenase.

### GMPR activity is dependent on pH and monovalent cations

The pH optimum of *E. coli* GMPR is about 8 (Mager & Magasanik, 1960), but considering the slightly acidic intracellular pH of mycobacteria during hypoxia (Rao *et al*., 2001; Vandal *et al*., 2009; Zhang *et al*., 1999b), we analyzed Msm and Mtb GMPR activities at pH values ranging from 6.2 to 9 (Fig. 2A). The pH optima of Msm GMPR and Mtb GMPR were 7.4 to 7.7 and 7.6 to 8.2, respectively. Mtb GMPR activity was, however, significantly lower than that of Msm GMPR at all pH values tested.

**Fig 2.**
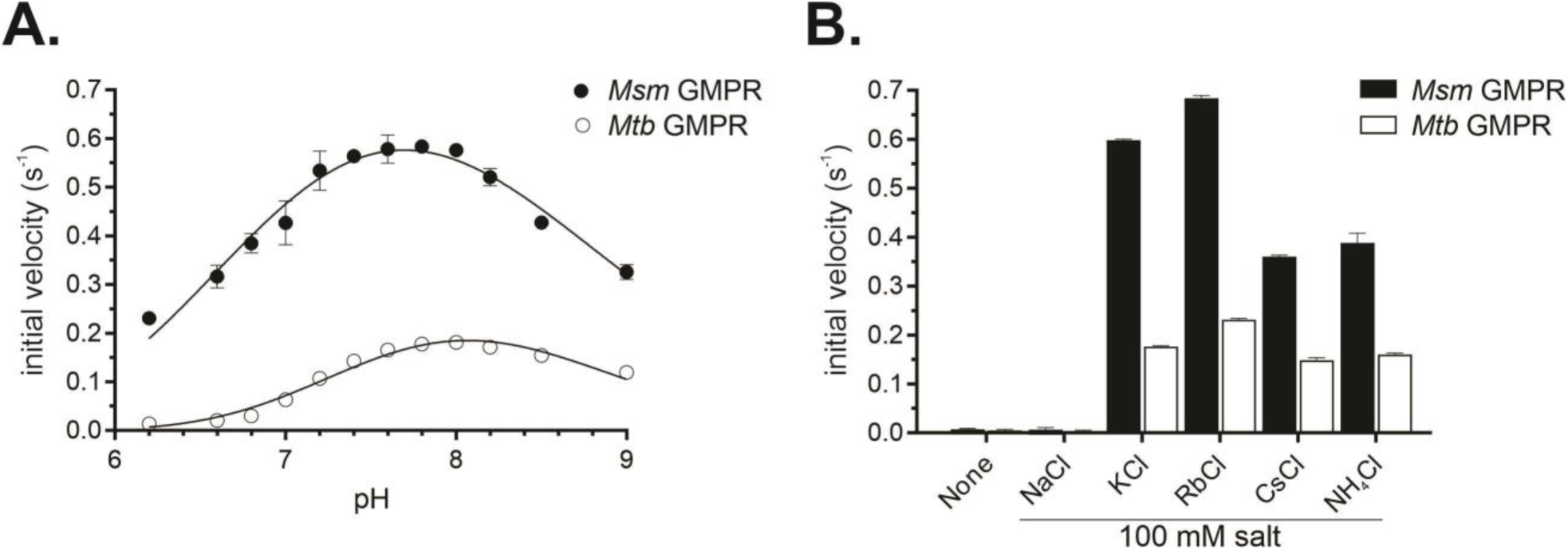
Dependence of Msm and Mtb GMPR activities on pH and monovalent cations. **A**) Determination of pH optima. Reaction velocities were measured in a mixture of 80 mM MPH buffer, 100 mM KCl, and 100 nM Msm GMPR at fixed concentrations of 200 μM NADPH and 100 μM GMP as substrates. **B)** Influence of monovalent cations on GMPR activity. Reaction velocities were measured at pH 7.6 in the presence of 100 mM corresponding salts, 200 μM NADPH, and 100 μM GMP.

Trypanosomal GMPRs require monovalent ions for catalytic activity; however, the effect of individual cations on these enzymes differs (Bessho *et al*., 2016; Sarwono *et al*., 2017). We analyzed the influence of monovalent cations on both Msm and Mtb GMPR activity and detected negligible activity in the presence of Na^+^ and increasing activity as follows: Cs^+^ ≍ NH_4_^+^ < K^+^ < Rb^+^ (Fig. 2B). For Msm GMPR, the initial velocities in the presence of Rb^+^ and K^+^ were 0.68 ± 0.02 s^-1^ and 0.60 ± 0.01 s^-1^, respectively. The reaction rates in the presence of Cs^+^ and NH_4_^+^ were reduced by approximately 40% to 0.36 ± 0.01 s^-1^ and 0.39 ± 0.03 s^-1^, respectively. Mtb GMPR exhibited a similar activity dependence on monovalent ions.

### GMPR activity is regulated by substrates and reaction products

Next, we measured the kinetic parameters of Msm and Mtb GMPR activity. The dependence of the Msm GMPR initial reaction velocity on the NADPH concentration at fixed GMP concentration (100 μM) followed Michaelis–Menten kinetics (Fig. 3A). The apparent Michaelis-Menten constant (K_m_) and apparent limiting initial velocity (V_lim_) values for NADPH were 30 ± 4 μM and 0.73 ± 0.03 s^-1^, respectively. On the other hand, the dependence of Msm GMPR initial reaction velocity on GMP concentration at a fixed NADPH concentration (200 μM) followed negative cooperative kinetics with a Hill coefficient (n_H_) of 0.53 ± 0.05 (Fig. 3B). The apparent K_0.5_ and V_lim_ values for GMP were 4.2 ± 0.6 μM and 0.66 ± 0.02 s^-1^, respectively. At pH 6.6, Msm GMPR activity significantly decreased (Table 1, Fig. S2). For Mtb GMPR, the apparent K_m_ value for NADPH at pH 7.8 was comparable with the apparent K_m_ value of Msm GMPR at pH 6.6, but the apparent V_lim_ value was 1.9 times lower (Table 1, Fig. S3). Like the Msm orthologue, Mtb GMPR showed negative cooperativity when velocity was plotted *versus* GMP concentration (Fig. S3).

**Fig 3.**
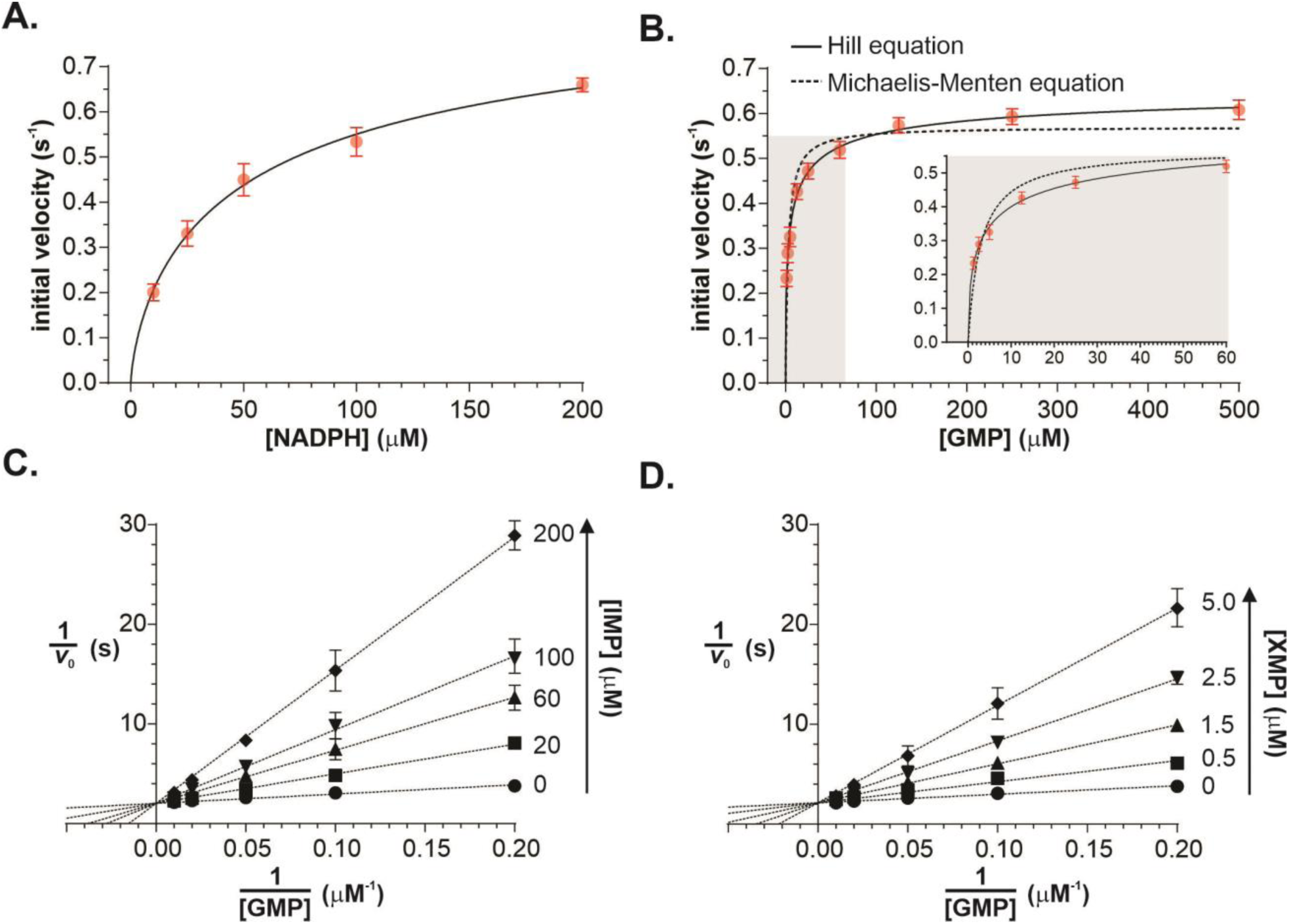
Steady state kinetics of the Msm GMPR-catalyzed reaction at pH 7.6. Reaction rates were measured in 80 mM MPH buffer, 100 mM KCl, and 20 nM Msm GMPR at pH 7.6 and 25 °C. **A)** Dependence of the initial reaction rate on the NADPH concentration at fixed GMP concentration (100 μM). The data were fitted with the Michaelis-Menten equation. **B)** Dependence of the initial reaction rate on the GMP concentration at fixed NADPH concentration (200 μM). Tha data were fitted with the Hill (solid line) or Michaelis-Menten (dotted line) equation. The inner graph is a magnification of the grey area. **C)** Inhibition of GuaB1 activity by IMP shown in Lineweaver-Burk representation. The IMP concentration is shown to the right of each line. **D)** Inhibition of Msm GMPR activity by XMP shown in Lineweaver-Burk representation. The XMP concentration is shown to the right of each line.

**Table 1:** Basic kinetic parameters of reactions catalyzed by Msm and Mtb GMPR under different conditions.

| pH | species | Varied<br>substrate /<br>ligand | $v_{lim}^{app2}$<br>$s^{-1}$ | $K_m^{app} (K_{0.5}^{app})$<br>μM | Dependence | $k_{cat}/K_m$<br>$\mu M^{-1} \cdot s^{-1}$ |
| --- | --- | --- | --- | --- | --- | --- |
| 7.6 | Msm | NADPH | $0.73 \pm 0.03$ | $30 \pm 4$ | MM eq. | 0.024 |
| 7.6 | Msm | GMP | $0.66 \pm 0.02$ | $4.2 \pm 0.6$ | Hill eq.,<br>$n_H 0.53 \pm 0.05$ | |
| 6.6 | Msm | NADPH | $0.51 \pm 0.01$ | $63 \pm 4$ | MM eq | 0.008 |
| 6.6 | Msm | GMP | $0.39 \pm 0.01$ | $3.5 \pm 0.3$ | Hill eq.,<br>$n_H 0.67 \pm 0.04$ | |
| 7.8 | Mtb | NADPH | $0.27 \pm 0.08$ | $63 \pm 7$ | MM eq. | 0.004 |
| 7.8 | Mtb | GMP | $0.13 \pm 0.01$ | $10.5 \pm 1.17$ | Hill eq.,<br>$n_H 0.52 \pm 0.03$ | |

Reaction products and substrate analogs often inhibit enzymatic reactions. Therefore, we evaluated Msm GMPR activity in the presence of IMP (product) and XMP (reaction intermediate). Our analysis showed that IMP and XMP are competitive inhibitors of Msm GMPR at pH 7.8 and 6.6 (Fig.3C, D, Fig. S2 B, C) with K_i_ values of 10 ± 0.4 μM and 0.6 ± 0.1 μM, respectively.

We also analyzed catalysis by Msm GMPR of the reverse reaction in the presence of 2 mM NADP^+^, 500 μM IMP, and 100 mM NH_4_Cl. However, under these conditions, we did not detect any measurable activity (data not shown).

### Msm GMPR activity is inhibited by ATP in a pH-dependent manner

We tested the effect of a physiologically relevant ATP or GTP concentration (1 mM) on Msm GMPR activity in the presence of 2 mM MgCl_2_ with fixed substrate concentrations. At pH 7.6, GTP had only a minor positive effect on the activity and ATP only a minor negative effect. At lower pH values, however, the effects of ATP and GTP increased (Fig. 4A). The inhibitory effect of 1 mM ATP increased sigmoidally with decreasing pH (7.8–6.4), plateauing at pH 6.6. In contrast, the activating effect of 1 mM GTP increased linearly with decreasing pH.

**Fig 4.**
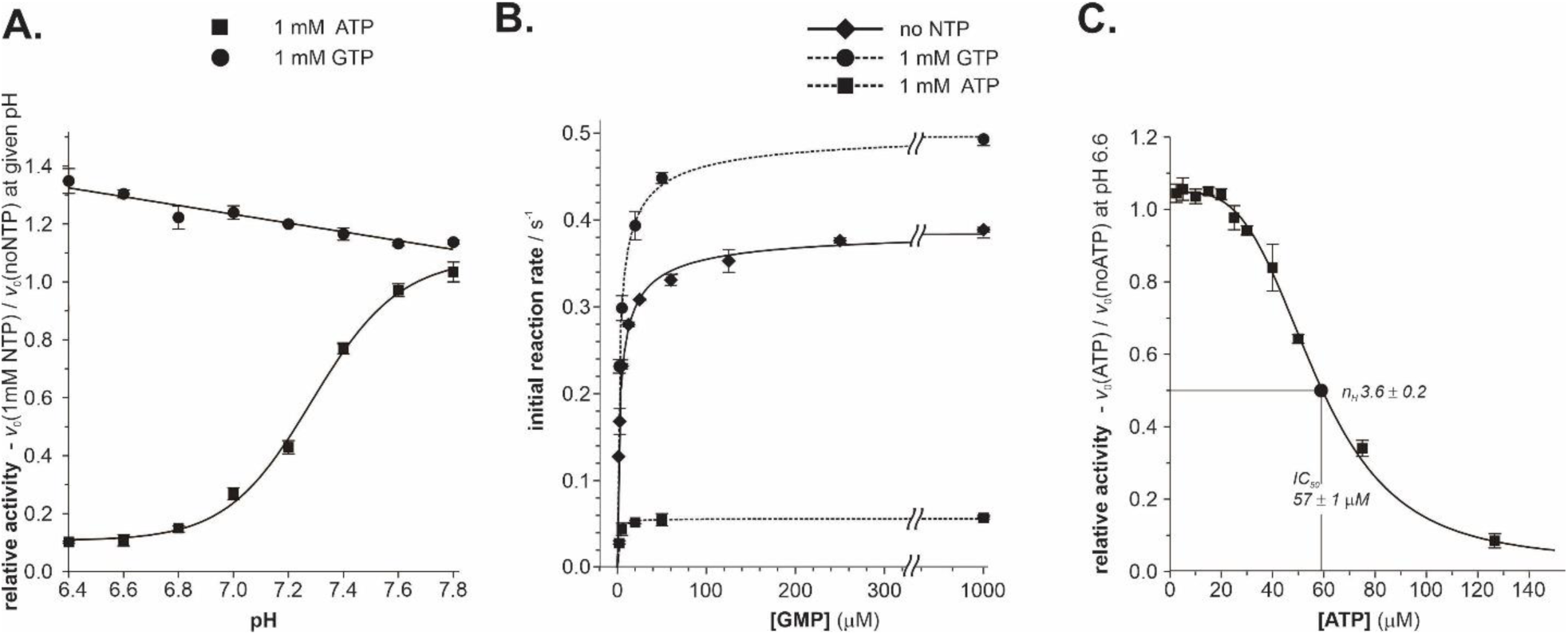
Influence of ATP and GTP on Msm GMPR activity. All reactions were performed at fixed substrate concentrations (100 μM GMP and 200 μM NADPH), 2 mM MgCl_2_, and 100 nM Msm GMPR in 80 mM MPH buffer at 25 °C. **A)** Dependence of the relative Msm GMPR activity on pH in the presence of 1 mM ATP or GTP. The relative activity is the ratio of the initial reaction velocity with 1 mM NTP and the velocity of the control reaction without NTPs. For ATP and GTP, the data were fitted with Hill and linear equations, respectively. **B)** Msm GMPR activity in the presence of 1 mM GTP and 1 mM ATP at pH 6.6 measured at a fixed 200 μM NADPH concentration. Plots show the initial rate *versus* GMP concentration. **C)** Dependence of the relative Msm GMPR activity on ATP concentration at pH 6.6. The relative activity is the ratio of the reaction rate at the given ATP concentration and the rate of the reaction without ATP. Data were fitted with the Hill equation.

To more deeply analyze the impact of GTP and ATP on GMPR activity at pH 6.6, we measured the initial velocity of reactions in the absence or presence of these nucleotides *versus* GMP concentration (Fig. 4B). At 1 mM concentrations, GTP and ATP caused only minor changes to n_H_ and K_0.5_. The n_H_ value changed from 0.67 ± 0.04 to 0.65 ± 0.06 and 1.29 ± 0.18, respectively, and the K_0.5_ value changed from 2.7 ± 0.3 to 3.5 ± 0.3 and 1.9 ± 0.1 μM, respectively. However, changes to V_lim_ were more substantial. Addition of 1 mM GTP increased the V_lim_ from 0.39 ± 0.01 to 0.51 ± 0.01 s^-1^ (a 36% increase), whereas addition of 1 mM ATP decreased V_lim_ to 0.056 ± 0.001 s^-1^ (an 86% decrease). Furthermore, we determined the dependence of the relative Msm GMPR activity on ATP concentration at pH 6.6 (Fig. 4C). The relative activity decreased sigmoidally with increasing ATP concentration; the IC_50_ value for ATP was 57 ± 1 μM.

### ATP-dependent Msm GMPR inhibition is restored by increasing GTP concentration

In our experimental setup, Msm GMPR activity is negatively regulated by ATP but positively regulated by GTP at slightly acidic pH. Both endpoint products of the purine metabolic pathway can concurrently affect Msm GMPR activity *in vivo*. We therefore measured Msm GMPR activity in the presence of 57 μM ATP (IC_50_) or 570 μM ATP (10×IC_50_) at increasing GTP concentrations (Fig. 5). At both ATP concentrations, the activity increased sigmoidally from maximally inhibited to fully recovered. In the presence of 57 μM ATP, half of the activity was recovered at 27 ± 2 μM GTP. In the presence of 570 μM ATP, half of the Msm GMPR activity was recovered at 470 ± 11 μM GTP. These results indicate that the activity of Msm GMPR can be tightly regulated by the ATP/GTP ratio.

**Fig 5.**
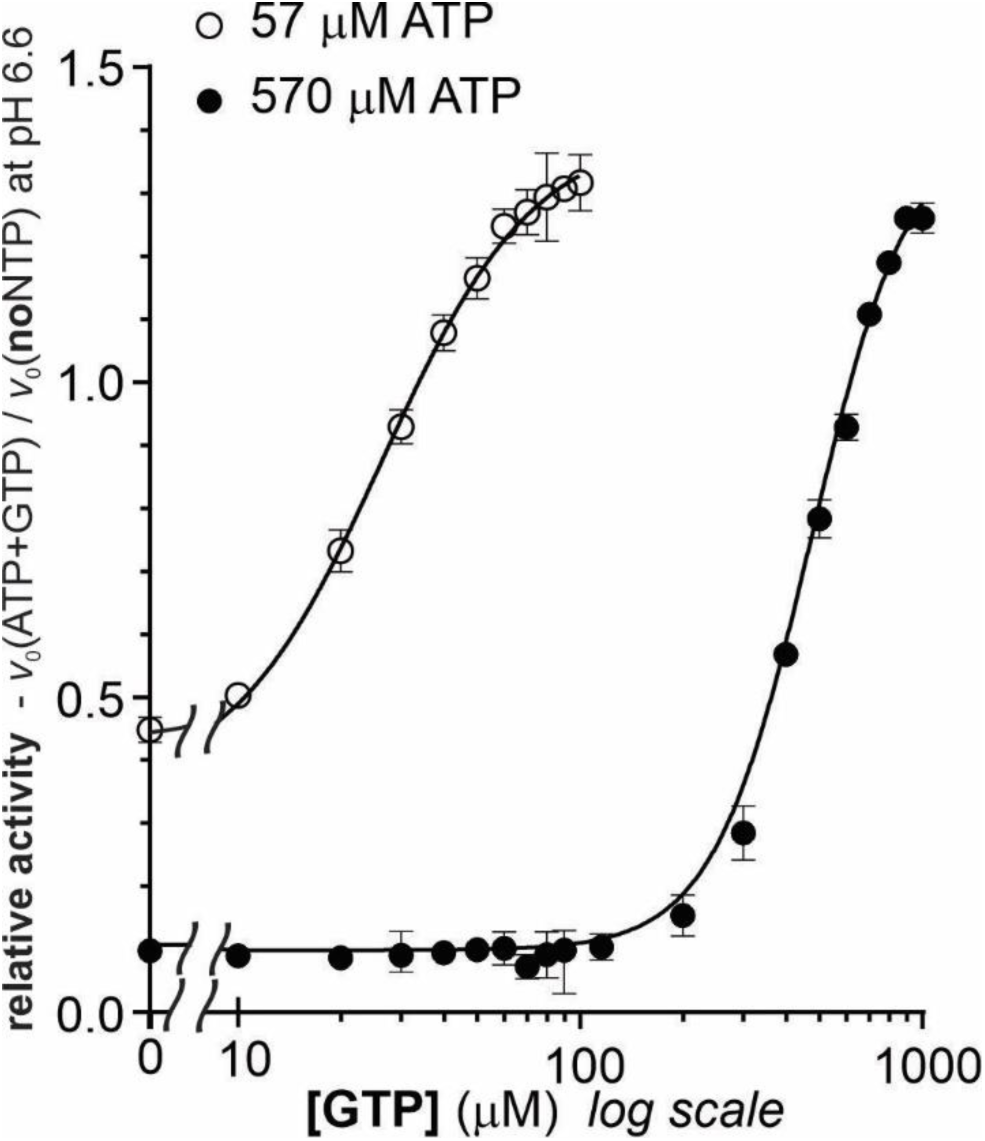
The effect of GTP on Msm GMPR activity inhibited by ATP. All reactions were performed at fixed substrate concentrations (100 μM GMP and 200 μM NADPH) and 100 nM Msm GMPR at 25 °C and pH 6.6. The activity of Msm GMPR at increasing concentrations of GTP was measured in the presence of 57 μM ATP (white circles) or 570 μM ATP (black circles). Relative activity is the ratio of the initial reaction velocities measured in the presence and absence of ATP and GTP.

### The oligomerization state of Msm GMPR is regulated by ligands and pH

To determine if the oligomerization state of Msm GMPR is regulated by nucleotide binding, we used size exclusion chromatography to analyze Msm GMPR in the presence of GMP, IMP, GTP, and ATP at various pH values (6.6, 7.3, 8.2). The oligomeric state of the protein was then expressed as the percentage of protein-forming tetramers (Fig. 6). In all experiments, we observed only tetramers or octamers.

**Fig 6.**
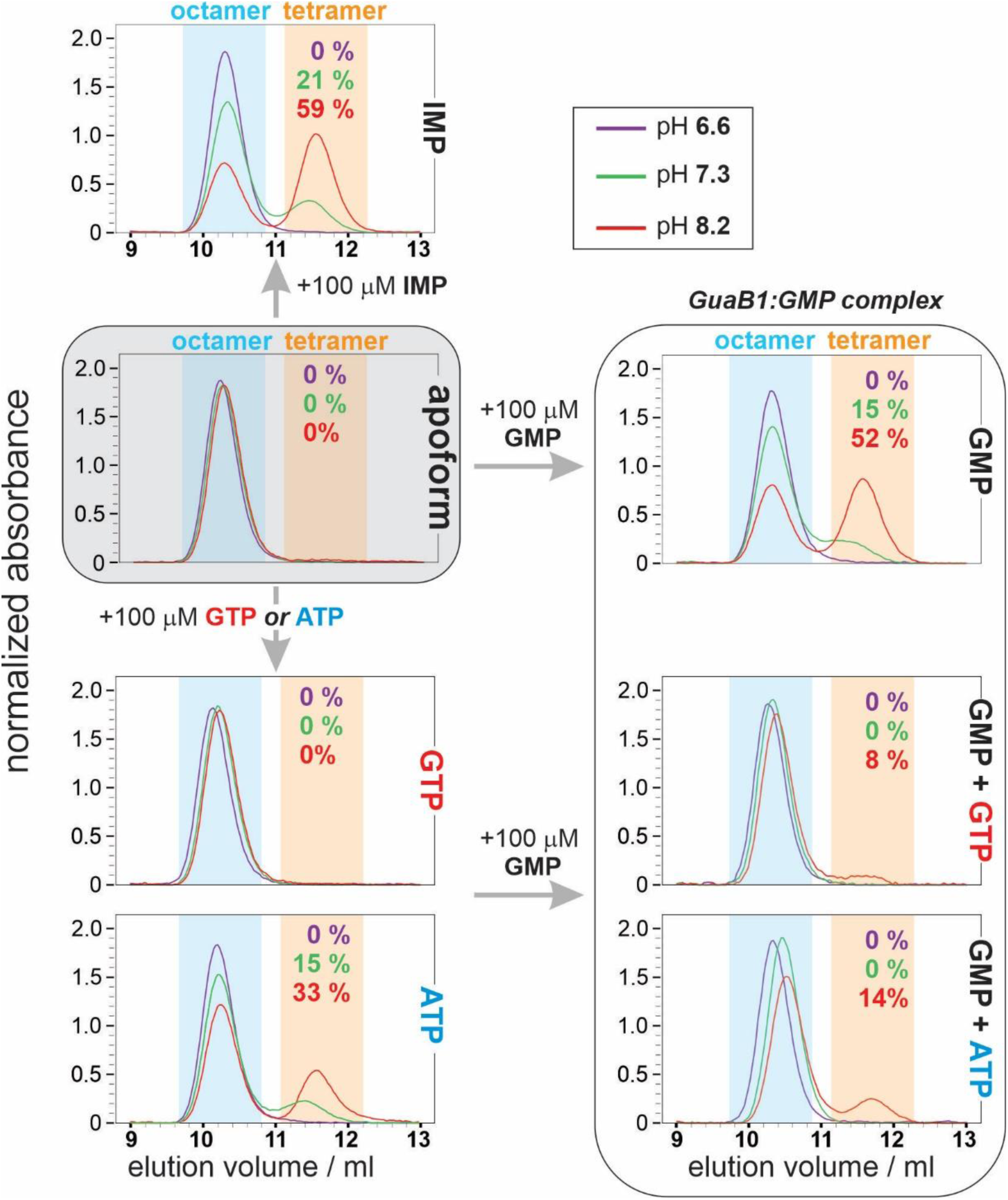
Analysis of Msm GMPR oligomerization. Msm GMPR (50 μM) was analyzed by size exclusion chromatography in the absence (apoform) or presence of 100 μM ligands at pH 6.6, 7.3, and 8.2. The oligomeric state of the protein is expressed as the percentage of protein forming tetramers.

The apoform of Msm GMPR and Msm GMPR with GTP were present only as octamers at all pH values tested. However, Msm GMPR with GMP, IMP, or ATP formed octamers and tetramers in a pH-dependent manner; the higher the pH, the stronger the effect of the ligands on dissociation of the octamers. At pH 8.2, 33–59% of the protein formed tetramers. At pH 7.3, 15– 21% of the protein formed tetramers. At pH 6.6, the effect of the ligands was not observable. This indicates that pH plays an important role in dissociation of Msm GMPR octamers to tetramers in the presence of a substrate (GMP), product (IMP), or activity effector (ATP).

We next analyzed the effect of GTP or ATP on the oligomeric state of the GMPR:GMP complex. Interestingly, both ligands induced octamerization of the complex. At pH 8.2, the percentage of Msm GMPR forming tetramers decreased from 52% to 8% or 14% when GTP or ATP was present, respectively. At pH 7.3 and 6.6, no tetramers were observed.

### X-ray crystallography reveals the unique position of CBS domains in the Msm GMPR octamer

*Trypanosoma brucei* GMPR is the only known GMPR structure that contains CBS domains (Imamura *et al*., 2020). The presence of CBS domains in GMPRs is unusual. However, analysis of the Msm GMPR amino acid sequence and its alignment with sequences of *E. coli* GMPR, which lacks a CBS domain, and Ld and Tb GMPRs, which have CBS domains, showed that residues S91–R214 in Msm GMPR are located at the position corresponding to the protozoal CBS (Fig. S4). To confirm the presence of a CBS domain in Msm GMPR and to learn how the structure of Msm GMPR differs from structures of other IMPDHs and GMPRs, we determined the structure of Msm GMPR by X-ray crystallography.

Msm GMPR crystallized in space group C2 with eight monomers in the asymmetric unit. The crystal parameters are listed in Table 2. The structure was deposited in the Protein Data Bank under PDB ID 7OY9. As expected, the Msm GMPR monomer is composed of two domains (Fig. 7A). The catalytic domain has a TIM barrel structure typical for the IMPDH/GMPR structural family (Fig. S5). The Msm GMPR sequence spanning residues 97– 207 folds as a typical Bateman domain composed of two CBS pairs. The Msm GMPR catalytic domains form a tetramer with a four-fold axis and the Bateman domains at the perimeter. Two tetramers interact through the Bateman domains, thus forming an octamer typical for IMPDHs and GPMRs with Bateman domains (Fig. 7B). However, unlike Tb GMPR (Fig. 7C) and other known structures of IMPDH and GPMR octamers in which the Bateman domains extrude from the octamer and thus have few contacts with the catalytic domains, the Bateman domains in Msm GMPR are very close to the catalytic domains and have a relatively large interacting area. The mutual orientation of the interacting Bateman domains in the Msm GMPR octamer is also unique. Usually, Bateman domains from opposite tetramers interact through both CBS domains, but the Msm GMPR Bateman domains have only a small interface (compare Fig. 7B and 7C).

**Fig 7.**
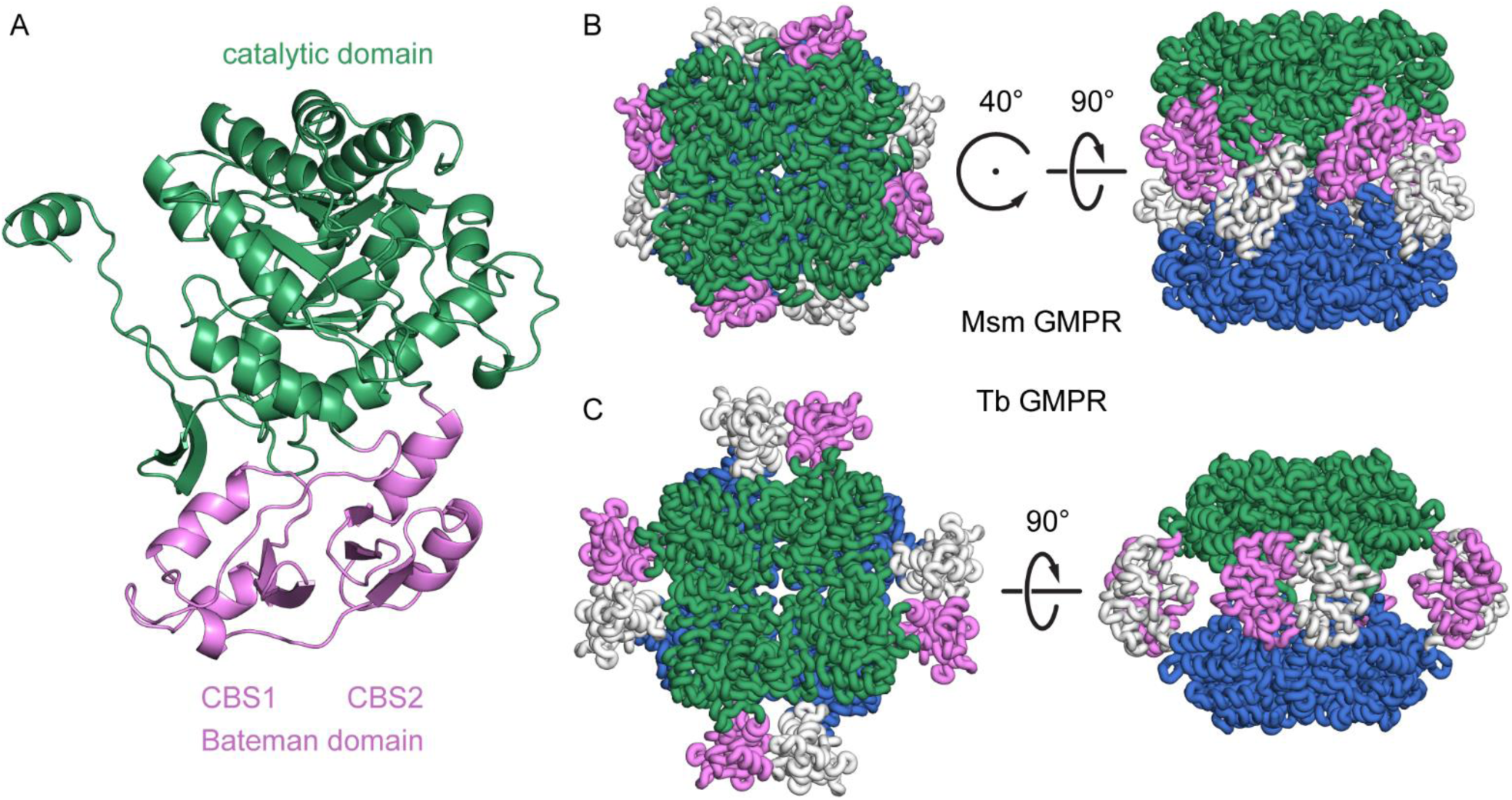
X-ray structure of Msm GMPR and comparison with the Tb GMPR structure. **(A)** The Msm GMPR monomer is composed of a catalytic domain with the TIM barrel structure typical of the IMPDH/GMPR family (green) and a Bateman domain composed of two CBS domains (pink). **(B)** Quaternary organization of the Msm GMPR octamer. **(C)** Quaternary organization of the Tb GMPR octamer (PDB ID: 6JL8). In both structures, the catalytic domains (green/blue) form a tetramer with four-fold symmetry, but the Bateman domains (pink/white) have different orientations and their interactions in GMPR are positioned at the perimeter.

### The CBS domain is necessary for Msm GMPR function *in vivo*

To test the importance of the CBS domain for GMPR function in Msm, we complemented the guanine-utilization-deficient Msm *ΔguaB1ΔpurF* strain with plasmids encoding Msm GMPR or Msm GMPR with deleted CBS domain (ΔCBSGMPR) (Fig. S6). Restored *ΔguaB1ΔpurF* growth on guanine as a sole purine supplement was observed only in the presence of full-length Msm GMPR. These results illustrate the indispensability of the CBS domain for Msm GMPR function. We did not succeed in purifying the ΔCBSGMPR mutant for complementary biochemical studies, although we prepared several constructs and used different purification procedures.

### The Msm guaB1 gene is an Actinobacteria phylum-specific feature

To determine whether GMPRs containing CBS domain homologues (GuaB1) are also present in other organisms, we searched for Msm GMPR homology protein sequences in bacterial genomes using BLAST. Our analysis revealed GuaB1 homologues throughout the entire phylum of *Actinobacteria* (Fig. 8). Genomes of *Actinobacteria* species contain two GuaB1 homologues with CBS domains. These GuaB1 sequence homologues typically contain two CBS domains inserted into a highly conserved IMPDH domain. Because of the principal sequence similarity between GMPRs and IMPDHs, we used the Hidden Markov Model (HMM) approach to confirm the identity of GuaB1 (Fig. S7). Genes for IMPDH in *Actinobacteria* are accompanied by another IMPDH homologue, designated as GuaB3 in mycobacteria (Rv3410c and MSMEG_1603 in Mtb and Msm, respectively).

**Fig 8.**
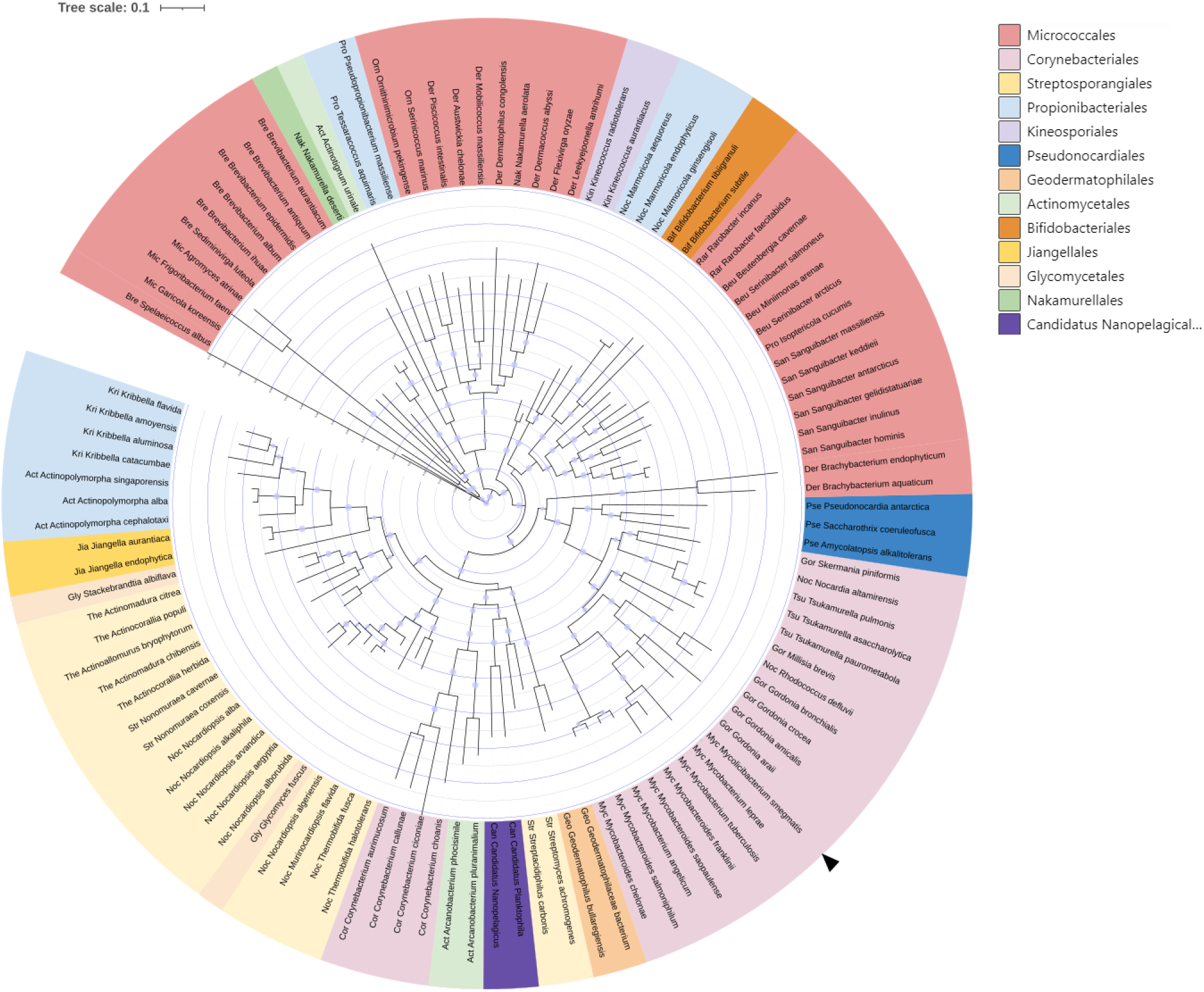
Phylogenetic tree constructed from GuaB1 homologues found throughout the *Actinobacteria* phylum. For clarity, the tree shows only a fraction of the organisms from the twelve orders. Orders are visually separated by color; family is indicated by the abbreviation in front of the organism’s name. Homologues were chosen to capture the diversity of the homologous sequence within the entire phylum and to show interrelationships. Blue dots represent the bootstrap value of replicate trees above 0.5.

## Discussion

The Msm and Mtb genomes each contain three *guaB* genes. Most studies have focused on Mtb *guaB2*, which encodes an essential IMPDH and is a potential drug target (Cox *et al*., 2016; Sahu *et al*., 2018; Singh *et al*., 2017; Usha *et al*., 2011). Two other genes, Mtb *guaB1* (Rv1843c) and *guaB3* (Rv3410c), and their Msm orthologues, Msm *guaB1* (MSMEG_1602) and *guaB3* (MSMEG_1603), have unknown function, although their products are annotated as members of the IMPDH family in databases.

Characterization of mutated Msm strains lacking *guaB1* and the essential *purF* gene involved in the *de novo* biosynthesis pathway showed that Msm GuaB1 is involved in guanine interconversion in the purine salvage pathway. Testing of *in vitro* activity and identification of reaction products using UPLC chromatography revealed that GuaB1 catalyzes NADPH-dependent conversion of GMP to IMP and thus serves as a 5’-monophosphate reductase. Results of positive complementation experiments in the Msm *ΔguaB1ΔpurF* strain with Msm and Mtb *guaB1* sequences, the high amino acid sequence similarity of Msm and Mtb *guaB1*-encoded proteins, and testing of recombinant Msm and Mtb GuaB1 activities demonstrated that *guaB1* encodes a GMPR in both Msm and Mtb.

The requirement of monovalent cations for activity is considered a characteristic feature of IMPDHs (Heyde *et al*, 1976; Kerr *et al*, 2000; Riera *et al*, 2011; Rostirolla *et al*, 2014; Xiang *et al*, 1996), while the majority of characterized GMPRs (*Homo sapiens*, *Salmonella typhymurium*, *Escherichia coli,* and *Mycoplasma mycoides*) do not need monovalent cations for their activities (Benson & Gots, 1975; Deng *et al*, 2002; Martinelli *et al*, 2011; Mitchell *et al*, 1978; Spector *et al*, 1979). Protozoal Tb GMPR and Tc GMPR display increased activity in the presence of K^+^ and NH_4_^+^ (Bessho *et al*., 2016; Sarwono *et al*., 2017), but the essentiality of ions for GMPR activity has not been demonstrated. Msm and Mtb GMPRs require alkali metal ions with the ionic radius of K^+^ or higher (Rb^+^, Cs^+^) for catalysis. Thus, to the best of our knowledge, mycobacterial GMPRs are the first GMPRs to be identified as strictly requiring monovalent ions for activity.

Biochemical characterization indicated that the kinetics of Msm GMPR-catalyzed reactions differ from those catalyzed by other GMPRs. GMPRs lacking a CBS domain follow Michaelis-Menten kinetics in dependence on the GMP concentration. Protozoal Lm GMPR and Tb GMPR, which contain CBS domains, follow positively cooperative kinetics in dependence on GMP (Imamura *et al*., 2020; Smith *et al*., 2016). However, Msm and Mtb GMPRs displayed negative cooperative kinetics dependent on the GMP concentration. Activity regulation by the end products of the purine biosynthesis pathway, ATP and GTP, is an important feature of protozoal and Msm GMPRs. In protozoal GMPRs, the presence of GTP switched the cooperative GMP kinetics to hyperbolic Michaelis-Menten kinetics (Imamura *et al*., 2020; Smith *et al*., 2016). Msm GMPR activity is also regulated by ATP and GTP, but in a pH dependent manner. At the optimal pH of 7.6, 1 mM GTP or ATP had only a minor effect on Msm GMPR activity. At lower pH, the positive and negative effects of GTP and ATP became apparent. At pH 6.6, 1 mM GTP caused a 31% activity increase and 1 mM ATP a 86% activity decrease. GTP can effectively diminish Msm GMPR inhibition by ATP. Addition of 1 mM GTP recovered full Msm GMPR inhibition by ATP and even increased it to the activity level in the presence of 1 mM GTP alone. Thus, the ratio of GTP and ATP concentrations can effectively regulate the activity of Msm GMPR, but only at slightly acidic pH.

Analysis of the oligomeric state of Msm GMPR by gel filtration revealed the roles of GMP (substrate), IMP (product), and ATP and GTP (activity effectors) on dissociation of octamers into tetramers at different pH values. The Msm GMPR apoform was octameric under all conditions tested. At pH 6.6, binding of substrate, product, or allosteric effectors did not lead to formation of tetramers; Msm GMPR was present in solution only as an octamer. However, at pH higher than 7, substrate or product binding led to formation of a tetrameric pool, which increased with increasing pH. Binding of ATP and GTP to the Msm GMPR-GMP complex shifted the octamer/tetramer ratio back to an octameric state. A low percentage of tetramers was detected only at pH 8.2. Thus, the intracellular pH significantly contributes to Msm GMPR activity and oligomeric state regulation by binding of allosteric effectors.

The unusual kinetics behaviour of Msm GMPR could be attributed to the presence of a CBS domain and its unusual orientation in the octamer. Msm GMPR contains an interspacing region in the catalytic domain, which forms a Bateman domain and is essential for enzyme activity, as documented by complementation experiments in the Δ*guaB1*Δ*purF* Msm strain with wt and Msm ΔCBS GMPR variants. The CBS domain is present exclusively in IMPDHs and protozoal GMPRs from *Leishmania major* and *Trypanosoma brucei*, and Msm GMPR is the first example of a bacterial GMPR with a CBS domain. The X-ray structure indicated that the Msm GMPR catalytic domain forms a tetramer with the Bateman domain at the perimeter, and two tetramers form an octamer through the CBS domains. However, the CBS domains are much closer to the catalytic domains and thus share a much larger interacting area in comparison with other known structures of IMPDHs. Importantly, construction of a phylogenetic tree based on Msm GuaB1 homologues indicated the presence of *guaB1-*encoded GMPR with two CBS domains inserted into a highly conserved catalytic domain throughout the *Actinobacteria* phylum.

Taken together, our data show that the Msm *guaB1* gene encodes a functional guanosine 5’-monophosphate reductase with a CBS domain and that its orthologues are spread across the *Actinobacteria* phylum and contributes to regulation of the purine nucleotide pool by recycling GMP to IMP.

## Material and methods

### DNA constructs

For all cloning procedures, we used Q5-polymerase (NEB) for PCR DNA amplification and *E. coli* DH5α for plasmid amplification. T4 DNA ligase and restriction endonucleases were purchased from NEB (USA). Ligation of DNA fragments was performed using In Fusion^TM^ cloning (Takara, Japan). Final constructs were verified by Sanger sequencing (EUROFINS, Germany). Primer sequences (Generi Biotech, Czech Republic) are listed in Supplementary Table 1.

Specific Msm *guaB1* and *guaB2* deletion cassettes (designated pYS2-Δ*guB1* or pYS2-Δ*guaB2*) were based on the pYS2 plasmid, which contains the *Spe*I/*Swa*I-*loxP-gfp-hyg^r^-loxP*-*Pac*I/*Nsi*I selection locus (Shenkerman *et al*, 2014). Regions upstream of *guaB1* (703 bp) and *guaB2* (889 bp) were amplified by PCR using Msm chromosomal DNA as a template with primer pairs 1/2 and 5/6, respectively, and ligated into pYS2 *via Spe*I/*Swa*I sites. Next, 829-bp *guaB1* and 747-bp *guaB2* downstream regions were amplified by PCR with primer pairs 3/4 and 7/8, respectively, and ligated into the corresponding pYS2-intermediate *via Pac*I/*Nsi*I sites. Construction of the deletion plasmid pYS2-Δ*purF* has been described previously (Knejzlik *et al*., 2019).

Msm GMPR complementation expression plasmids were constructed based on the pSE200 vector, derived from the pSE100 plasmid containing a constitutive Pmyc1TetO promoter in the absence of a Tet repressor (Ehrt *et al*, 2005). The pSE200-Msm.GuaB1 and pSE200-Mtb.GuaB1 plasmids were constructed as follows: *Msm.guaB1* and *Mtb.guaB1* were amplified from genomic DNA by PCR using primer pairs 9/10 and 11/12, respectively, and the fragments were inserted into *Swa*I-linearized pSE200 by the In-Fusion approach. Plasmid pSE200-Mtb.GuaB1.His for expression of the C-terminally His-tagged Mtb GMPR in Msm was constructed similarly using primer pair 23/24. The pSE200-Msm.GuaB1Δ(S91-R214) plasmid was constructed as follows: the pSE200-Msm.GuaB1 plasmid was PCR amplified using primer pair 13/14, and the PCR product was circularized by the In-Fusion approach. The pTriex-Msm.GuaB1 plasmid was constructed as follows: the *Msm guaB1* gene was amplified by PCR by primer pair 17/18 and inserted into the PCR-linearized pTriex-4 vector (Novagen, USA) with primers 15/16 by the In-Fusion approach. The plasmid for expression of the His.S91-R214 Msm GMPR fragment was constructed as follows: the corresponding sequence region was amplified by PCR using primer pair 21/22 and ligated *via* the In-Fusion approach to PCR-linearized pTriex-4 by primer pair 19/20.

### Cultivation of M. smegmatis

The *M. smegmatis* mc^2^ 155 strain and its mutants were propagated in liquid 7H9 medium (Sigma) with 10% ADC supplement (5% BSA, 0.85% NaCl, 2% dextrose) and 0.05% tyloxapol or 7H10-ADC agar medium (Sigma) at 37 °C. In both cases, the carbon source was enriched with 0.5% (vol/vol) glycerol. Hygromycin and kanamycin were added at final concentrations of 150 and 25 μg/ml, respectively. Purine supplements (adenine, guanine and hypoxanthine) were prepared as 75 mM stocks in DMSO and were added to a final concentration of 200 μM when required.

### Gene deletion

Gene disruption using a pYS2 deletion plasmid was performed as previously described (Shenkerman *et al*., 2014). Briefly, the *Spe*I/*Nsi*I-linearized pYS2-Δ*guaB1* or pYS2-Δ*guaB2* deletion cassette was introduced into electrocompetent *M. smegmatis* cells with 0.2% acetamide-driven expressed Chec9 DNA by electroporation. Msm recombinants were obtained by selection on 7H10/ADC medium containing hygromycin at 42 °C for 3 days. In the case of *guaB2* gene, the medium was supplemented with 200 μM guanine. To remove the Hyg^r^ cassette, gene disruptants were transformed with the pML2714 vector, which constitutively expresses Cre recombinase. Deletions were screened by PCR using Q5 polymerase and primer pairs that anneal at the boundaries of the deleted regions, and amplicons were sequenced. In the Δ*guaB2* strain, the second deletion of the *purF* gene was not cleared from the Hyg^r^ cassette.

### Nucleotide ligands

Sodium salts of GMP, IMP, XMP and NADPH were purchased from Santa Cruz. Sodium salts of ATP and GTP were from Sigma. Except for NADPH, the compounds were dissolved in deionized water to final concentrations of 50–100 mM. The pH of the ATP and GTP stocks was immediately adjusted to pH 7.0 with 1 M NaOH. The exact concentrations of the stocks were determined spectrophotometrically at the absorption maxima using the following absorption coefficients: ε^260 nm^ (ATP) 15.4 mM^-1^cm^-1^, ε^253 nm^ (GTP, GMP) 13.7 mM^-1^cm^-1^, ε^263 nm^ (XMP) 8.6 mM^-1^cm^-1^, and ε^249 nm^ (IMP) 12.1 mM^-1^cm^-1^. The stocks were aliquoted and stored at −70 °C. NADPH was dissolved in deionized water directly before measurements to create a 20 mM solution. The exact concentration was determined spectrophotometrically at 340 nm using the extinction coefficient ε^340 nm^ (NADPH) 6.3 mM^-1^cm^-1^.

### Protein expression and purification

Recombinant Msm GMPR and Msm CBS (S91–R214 fragment) with uncleavable C-terminal and N-terminal histidine tags, respectively, were produced in *Escherichia coli* BL21(DE3) RIL cells. ZYM-505 medium (Studier, 2014) containing ampicillin (50 µg/ml) was inoculated with an overnight culture of cells carrying the appropriate expression plasmid to an initial OD_590_ of 0.1. The resulting culture was cultivated at 37 °C to an OD_590_ of 2.0–3.0. The temperature then was lowered to 18 °C and expression was induced by 0.4 mM IPTG. The cells were harvested 16 h after induction at 8,000 x *g* for 10 min. The cell pellet was resuspended in lysis buffer (200 mM potassium phosphate, pH 8.0, 2 M potassium chloride, 2.5 mM TCEP, 10 mM imidazole, 0.5% Triton X-100, 1 mM PMSF, 0.1 mg/ml lysozyme) and stirred for 60 min at 4 °C. The lysate was sonicated, and the insoluble fraction was removed by centrifugation at 35,000 x *g* for 30 min. The supernatant was loaded onto an immobilized metal affinity chromatography column (HiTrap IMAC HP 5 ml) charged with Ni^2+^ and equilibrated with buffer A (200 mM potassium phosphate, pH 8.0, 2 M potassium chloride, 2.5 mM TCEP) containing 10 mM imidazole. The column was washed with buffer A containing 82.5 mM imidazole, and the His-tagged proteins were eluted with buffer A containing 300 mM imidazole. Msm GMPR and Msm CBS were next purified by size exclusion chromatography using HiLoad 26/60 Superdex 200 pg and HiLoad 26/60 Superdex 75 pg columns in buffer A, respectively. The purified proteins were transferred into storage buffer (50 mM Tris, pH 8.0, 2.5 mM TCEP) by desalting (HiPrep 26/10 Desalting) and concentrated to 25 mg/ml by centrifugal ultrafiltration. Finally, the proteins were aliquoted and stored at −70 °C. All purification procedures were performed on ice or at 4 °C. All FPLC equipment was manufactured by GE Healthcare Life Sciences. The purity of the target proteins was analyzed by SDS-PAGE, and the identity of the proteins was confirmed by mass spectrometry. The concentration and identity of copurified nucleotide 5’-monophosphates were determined by HPLC.

Recombinant Mtb GMPR with an uncleavable C-terminal His tag was produced in Msm with a deleted *guaB1* gene (*ΔguaB1*) under the control of the constitutive Psmyc promoter. The Msm *ΔguaB1* strain tranformed with plasmid pSE200-Mtb.GuaB1.His was cultivated in LB medium containing 0.1% (w/v) tyloxapol and 0.5% (w/v) glycerol until the OD_590_ reached 10. The cell pellet from 5 l media was resuspended in 200 ml lysis buffer and disintegrated by 3 passes on a French press at a pressure of 1,500 psi. The crude lysate was centrifuged at 50,000 x *g* for 40 min. The supernatant was loaded onto an immobilized metal affinity chromatography column (HiTrap IMAC HP 5 ml) charged with Ni^2+^ and equilibrated with buffer A containing 50 mM imidazole. The column was washed with buffer A containing 100 mM imidazole, and the His-tagged Mtb GMPR was eluted with buffer A containing 300 mM imidazole. Purified Mtb GMPR was desalted into storage buffer using a PD10 column (GE Healthcare), concentrated to 2 mg/ml by centrifugal ultrafiltration, aliquoted, and stored at −70 °C. All purification steps were carried out at 4 °C.

### Enzyme kinetics

All enzymatic reactions were carried out in MPH buffer composed of 30 mM MES (2-(N-morpholino) ethanesulfonic acid), 30 mM HEPES (2-[4-(2-hydroxyethyl) piperazin-1-yl]ethanesulfonic acid), 20 mM PIPES (2-[4-(2-sulfoethyl)piperazin-1-yl]ethanesulfonic acid), 100 mM KCl and 2 mM DTT and adjusted to the appropriate pH with 10 M NaOH. Msm GMPR was used in the concentration range 20–100 nM; actual concentrations are stated for individual experiments. Msm GMPR did not show any deviations in specific activity in this concentration range. The data were processed with GraphPad Prism 7 software.

The initial reaction velocity of the GMPR reaction was determined from the decrease in NADPH absorbance at 340 nm during the reaction. The absorbance was measured in 20 s intervals for 30–60 min in a 1 cm quartz cuvette at 25 ± 0.1 °C using a Specord 200 PLUS spectrophotometer (Analytik Jena, Germany). The slope of the decrease was calculated from the linear part of the steady-state of the absorbance curve by least-squares linear regression. The initial reaction rate (s^−1^) was then calculated from equation 1

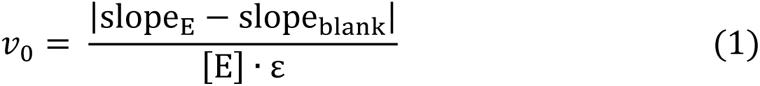

where slope_E_ and slope_blank_ are the slopes of the absorbance decrease at (A·s^−1^) for a reaction mixture containing Msm GMPR, NADPH, and GMP or for a blank mixture containing only Msm GMPR and a corresponding concentration of NADPH at the same conditions (pH, ionic strength, etc.), respectively. [E] is the molar concentration of Msm GMPR in nM and ε is the absorption coefficient of NADPH at 340 nm in nM^-1^cm^-1^ units (6.22·10^−6^).

The kinetic parameters of Msm GMPR were determined at pH 7.6 and 6.6. A reaction mixture containing NADPH and GMP in MPH buffer was equilibrated to 25 °C, and the reaction was started by addition of 20 nM Msm GMPR. The parameters of the reaction at a fixed concentration of GMP (100 μM) were calculated from the initial reaction rates for concentrations of NADPH in the range of 10–200 μM using equation 2.

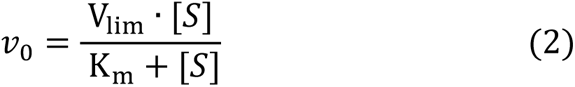

The parameters of the reaction at a fixed concentration of NADPH (200 μM) were calculated from the initial reaction rates for concentrations of GMP in the range of 1–500 μM using

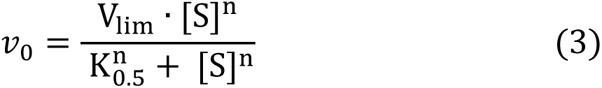

The inhibition constants for XMP and IMP at a fixed concentration of NADPH (200 μM) were calculated from the initial reaction rates for concentrations of GMP in the range of 5–100 μM using equation 4. Parameters K_i_, V_lim_ and K_m_ were shared for all IMP (0–200 μM) and XMP (0– 5 μM) concentrations.

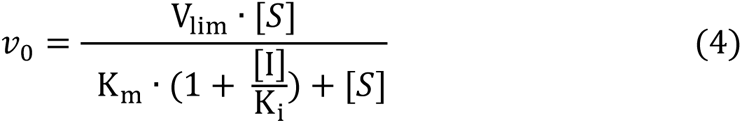

The effect of pH on the activity of Msm GMPR in the presence of ATP or GTP was measured as follows: 250 μl of 200 nM Msm GMPR in MPH buffer containing 2 mM MgCl_2_ at the given pH was preincubated without a ligand or with 1 mM GTP or ATP for 30 min at 25 °C. The reaction was started by addition of 250 μl preheated two-fold concentrated substrate mix containing 200 μM GMP and 400 μM NADPH in the same buffer (including 1 mM GTP or ATP as appropriate). The activity at the given pH was expressed as relative activity calculated from equation 5,

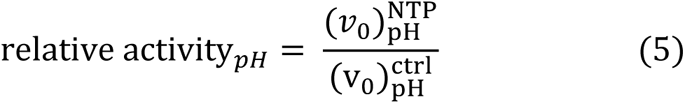

where 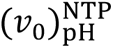 and 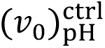 are initial velocities of the reaction at the given pH in the presence or absence of the ligand, respectively. The relative activities were plotted against pH and fitted with a linear (GTP) or Hill equation (ATP).

The effect of ATP concentration (1–125 μM) on Msm GMPR activity at pH 6.6 was measured as follows: 250 μl of 200 nM Msm GMPR in MPH buffer containing 2 mM MgCl_2_ was preincubated with ATP for 30 min at 25 °C. The reaction was started by addition of 250 μl preheated two-fold concentrated substrate mix (200 μM GMP and 400 μM NADPH) with the same ATP concentration. The activity at the given ATP concentration was expressed as relative activity calculated from equation 6,

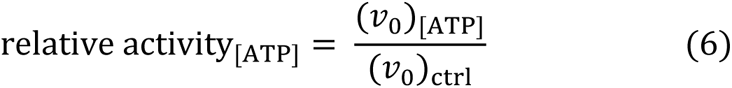

where (*v*_0_)_[ATP]_ and (*v*_0_)_ctrl_ are initial rates of a reaction containing ATP at a given concentration and a control reaction without ATP, respectively. The relative activity was plotted against ATP concentration and fitted with the Hill equation.

The ability of GTP to restore partially or fully ATP-inhibited Msm GMPR at pH 6.6 was measured as follows: 250 μl 200 nM Msm GMPR in MPH buffer containing 2 mM MgCl_2_ was preincubated with 57 μM (partial inhibition) or 570 μM ATP (complete inhibition) for 30 min at 25 °C. The reaction was started by addition of 250 μl preheated two-fold concentrated substrate mix (200 μM GMP, 400 μM NADPH) with ATP (57 or 570 μM) and GTP (two-fold final concentration). The activity at the given GTP concentration was expressed as relative activity calculated from equation 7,

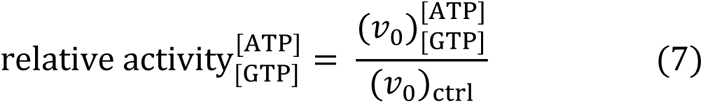

where 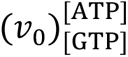 is the initial rate of a reaction containing ATP (57 or 570 μM) and GTP (0–1 mM), and (*v*_0_)_ctrl_ is the initial rate of a control reaction without ATP or GTP. The relative activity was plotted against GTP concentration and fitted with the Hill equation.

### Chromatographic analysis of the GMPR reaction mixture

Mixtures containing 75 μM NADPH and 100 μM GMP with or without 50 nM purified GMPR in MPH buffer (pH 7.6) in a total volume of 500 μl were incubated at 25 °C for 30 min. To remove the protein, 200 μl of the mixture was passed through a minispin 3 kDa cut-off centricon (Amicon, USA). The filtrate was analyzed by UPLC using an ACQUITY HSS T3 column (Waters, USA) equilibrated with a mobile phase composed of 50 mM potassium phosphate (pH 3.1) and 3 mM tetrabutylammonium bisulphite at a flow rate of 0.4 ml/min. After loading the sample, a linear gradient of 5–30% acetonitrile per 10 min was applied. The elution was monitored spectrophotometrically at 200–360 nm using a diode array detector. The compounds were assigned to individual elution peaks based on 50 μM calibration standards for NADPH, NADP^+^, GMP and IMP.

### Analytical size exclusion chromatography

The influence of selected ligands on the oligomeric state of Msm GMPR was analyzed by size exclusion chromatography (Superdex 200 Increase 10/300 GL, 1 ml/min, 25 °C). Frozen aliquots of the concentrated protein were diluted with running buffer (45 mM MES, 30 mM PIPES, 45 mM HEPES, 100 mM KCl, 0.5 mM TCEP, 2 mM MgCl_2_) containing appropriate ligands. The final concentration of the loaded protein was 2.5 mg/ml and the final volume was 10 µl. Protein elution was monitored by UV absorbance at 280 or 295 nm with respect to the absorption properties of the ligand. The chromatographic data were processed with Fityk software. The data were normalized and then fitted with an exponentially modified Gaussian function. The oligomeric state of the protein was expressed as the percentage of the protein forming tetramers.

### Protein crystallization

The initial conditions for MsmGMPR crystallization were screened with Morpheus (Molecular Dimensions) and JSCG Core I Suite (QIAGEN) protein crystallization screening kits.

Crystals were grown in a mixture of 0.3 μl protein solution and 0.3 μl reservoir solution in sitting drops at 19 °C in 96-well plates. The crystallization trials were set up with a Mosquito crystallization workstation (SPT Labtech). Initial conditions were further optimized by changing the protein concentration, buffer pH and precipitant concentration. The final crystals were grown in drops containing 22 mg/ml MsmGMPR, 0.03 M MgCl_2_, 0.03 M CaCl_2_, 20% ethylene glycol, 10% PEG 8000, 0.1 M Tris/bicine, pH 8.3. The harvested crystals were flash-frozen in liquid nitrogen.

### Data collection and structure determination

The data were collected at the MX14.2 beamline at BESSY, Berlin, Germany (Mueller *et al*, 2015), and processed using XDS (Kabsch, 2010) with XDSApp GUI (Krug *et al*, 2012). The initial structure was obtained by molecular replacement using the structure of inosine monophosphate dehydrogenase catalytic domain from *Streptococcus pyogenes* (PDB ID 1ZFJ) as a model (Zhang *et al*, 1999a). The initial structure was then improved by iterative manual rebuilding in Coot (Emsley *et al*, 2010) and automatic refinement in Phenix.refine (Afonine *et al*, 2012; Liebschner *et al*, 2019). The Bateman domains were build manually in the process. The final refined structure was validated with MolProbity (Williams *et al*, 2018). Data collection and refinement statistics are summarized in Table 2.

### GuaB1 tree reconstruction and consensus sequence

Sequences for phylogenetic analysis were retrieved from the NCBI non-redundant protein sequences(nr) database using the similarity search program blastp (similarity matrix BLOSSUM 62)(Altschul *et al*, 1990). The domain composition of the sequences was obtained from the Conserved Domain Database (CDD) (Yang *et al*, 2020) and served to confirm the identity of GuaB1. The sequences were filtered, redundant sequences were removed, and the number of sequences used to construct the phylogenetic tree was reduced by random selection to maintain the diversity representing the different orders of actinobacteria. The analysis was performed on the Phylogeny.fr platform (Dereeper *et al*, 2008). Sequences were aligned with MUSCLE (v3.8.31) (Edgar, 2004). Ambiguous regions were removed with Gblocks (v0.91b) (Castresana, 2000) with settings as follows: no gaps allowed in the final alignment, minimum number of sequences for a flank position: 85%; contiguous non-conserved positions larger than 8 were removed. The phylogenetic tree was reconstructed using the maximum likelihood method (Guindon & Gascuel, 2003). The gamma parameter was estimated from the data (gamma=0.733). Internal branch reliability was assessed by the aLRT test (SH-Like). The phylogeny tree was visualized with the iTOL platfrom (Letunic & Bork, 2021). The domain composition of the consensus sequence was explored using InterProScan (Jones *et al*, 2014). The consensus logo and visualization were created in Jalview (v2.11.1.4) (Anisimova & Gascuel, 2006).

## ACKNOWLEDGEMENT

We thank Dagmar Grundova for excellent technical support. This work was supported by the European Regional Development Fund; OP RDE; Project No.

CZ.02.1.01/0.0/0.0/16_019/0000729 and by RVO project 61388963.

## AUTHOR CONTRIBUTIONS

Conceptualization, Z.K., M.D. and I.P.; Investigation, Z.K., M.D., K.C., M.K., M.D., E.Z. and D.R.; Formal analysis, Z.K., M.D., K.C., M.K. and E.Z; Writing – original draft, Z.K. and M.D.; Writing – review & editing, I.P.; Funding acquisition, I.P.

## DECLARATION OF INTERESTS

The authors declare no competing interests.

## Supplementary Method

### Metabolite analysis

7H9/ADC medium (100 ml in a 500 ml Erlenmeyer cultivation flask) was inoculated with a fresh culture of Msm at the stationary phase (OD_600_ = 10) to an initial OD_600_ of 10^-3^. The cell suspension was grown to an OD_600_ of 0.5 at 37 °C and 200 rpm (15–18 h). Then, 3 ml of the cell suspension was quickly vacuum-filtered through a 0.45 μm/25 mm cellulose acetate filter. The membrane with the collected bacteria was immediately transferred into 1 ml ice-cold 1 M acetic acid in a 1.5 ml microtube and quickly frozen in liquid nitrogen. The samples were slowly thawed on ice and then incubated on the ice for 30 min with short vortexing at 5 min intervals. The crude bacterial lysate was separated from the filter by centrifugation (4000×*g*, 30 s) through a pinhole at the bottom of the 1.5 ml microtube inserted into a 2 ml collection tube. The lysate was flash-frozen in liquid nitrogen and lyophilized. The material was then resuspended in 200 μl ice-cold deionized water and incubated on ice for 30 min. Insoluble material was removed by centrifugation at 22,000×*g* for 20 min at 4 °C. Clear aqueous bacterial extract was collected and stored at −80 °C for subsequent analysis. Analysis of Msm extracts was performed on cZIC-HILIC columns (150 × 2.1 mm) with a flow rate of 0.3 mL/min. Mobile phase A was 10 mM ammonium acetate, pH 5, adjusted with acetic acid. Mobile phase B was 10 mM ammonium acetate, pH 5, and 90% acetonitrile. MS quantification was performed by negative electrospray ionization with the following parameters: capillary voltage = 2 kV, cone voltage = 20 V, source temperature = 120 °C, desolvation temperature = 400 °C, desolvation gas flow = 400 L/h, cone gas flow = 30 L/h.

The intracelular concentrations of metabolites were calculated from equation 12, where _[*metabolite*]*ex*_ is the concentration of the metabolite in the extract in μM, _*Vex*_ is the volume of the extract (μl), OD is the optical density of the cell suspension, and _*Vc*_ is the volume of the filtered cell suspension (ml). NC (2.3×10^7^ ml^-1^) is a number of cells in 1 ml of cell suspension at OD_600_ = 0.1 and was determined by a colony-forming unit assay. V_cell_ (5.2×10^-9^ μl) is the volume of a rod-shaped mycobacterial cell of the average dimensions 5.5×1.1 μm.

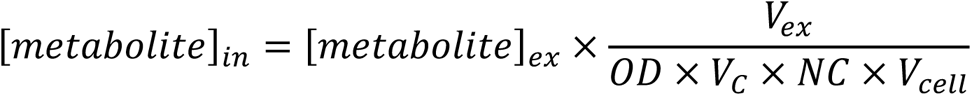

## Supplementary Figures

**Fig. S1:**
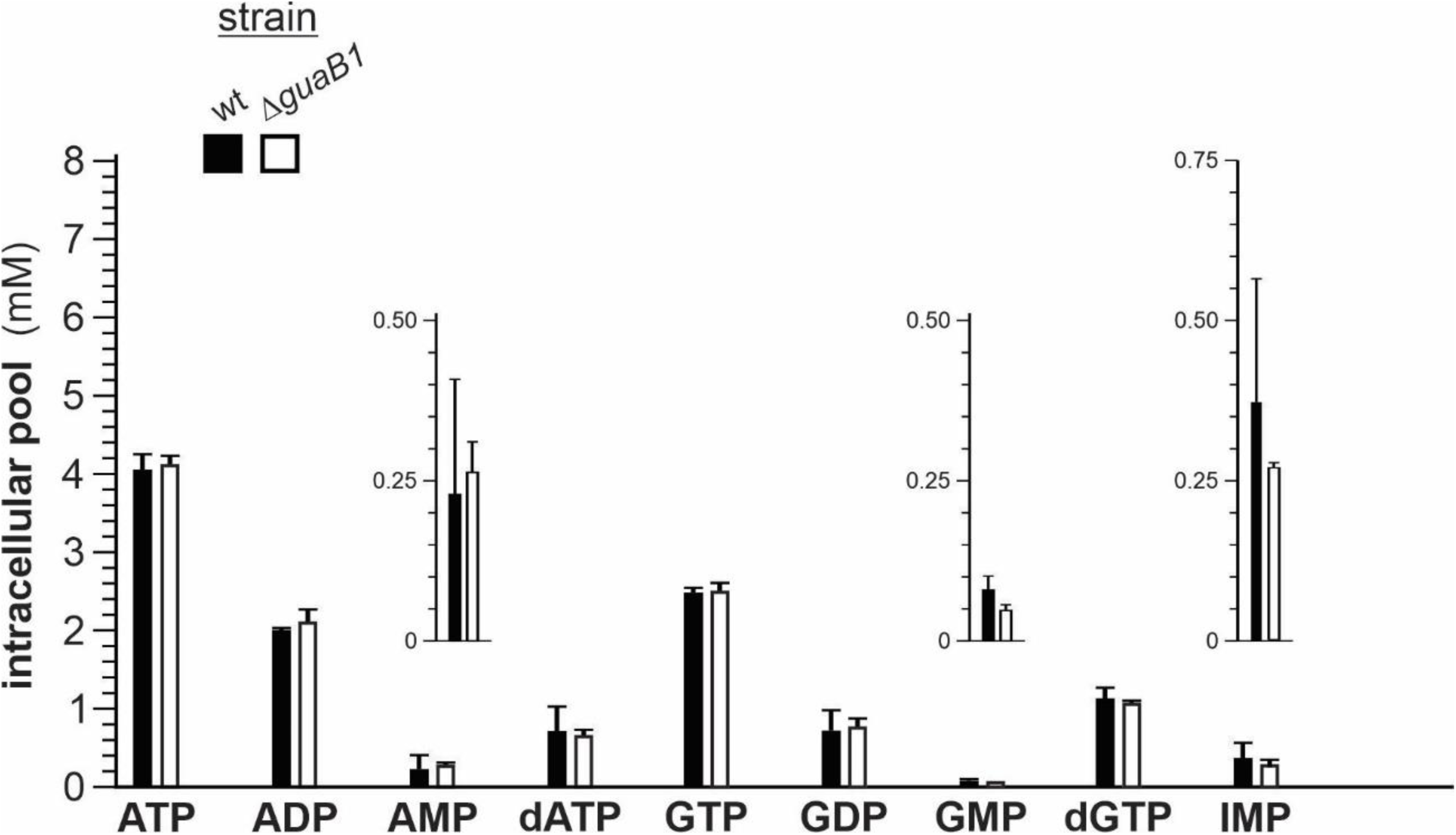
Intracellular purine metabolite analysis in wt and *ΔguaB1* Msm strains. Exponentially growing cells were collected by filtration and quickly lysed with 1 M acetic acid. The extracted nucleotides were analyzed by HILIC chromatography coupled with MS spectrometry and UV detection.

**Fig. S2.**
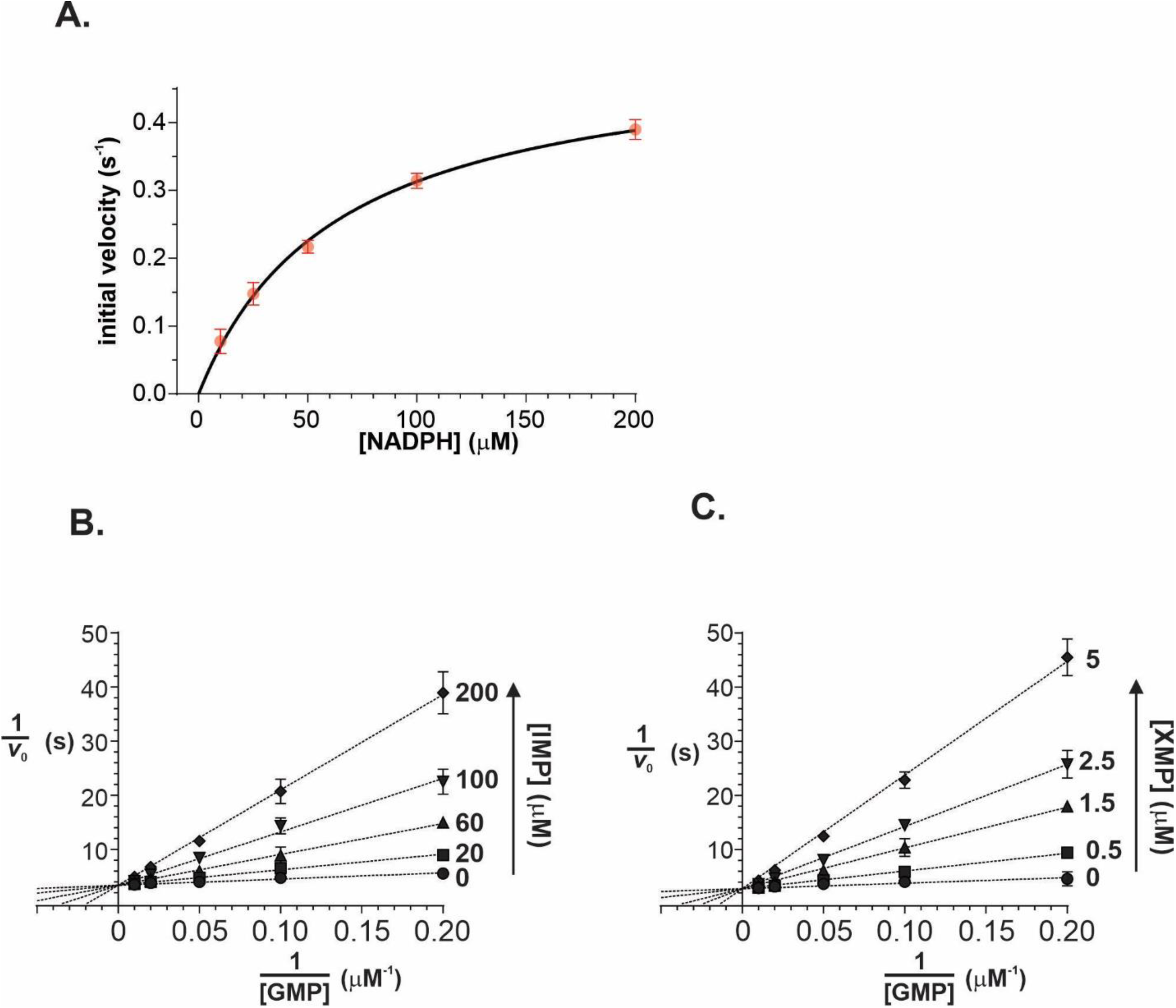
Steady state kinetics of the Msm GMPR-catalyzed reaction at pH 6.6. Reaction rates were measured in 80 mM MPH buffer, 100 mM KCl and 20 nM Msm GMPR at pH 6.6 and 25 °C. **A)** Dependence of the initial reaction rate on the NADPH concentration at fixed GMP concentration (100 μM). The data were fitted with the Michaelis Menten equation. **B)** Inhibition of GuaB1 activity by IMP shown in Lineweaver–Burk representation. The IMP concentration is shown to the right of each line. **C)** Inhibition of Msm GMPR activity by XMP shown in Lineweaver–Burk representation. The XMP concentration is shown to the right of each line.

**Fig. S3.**
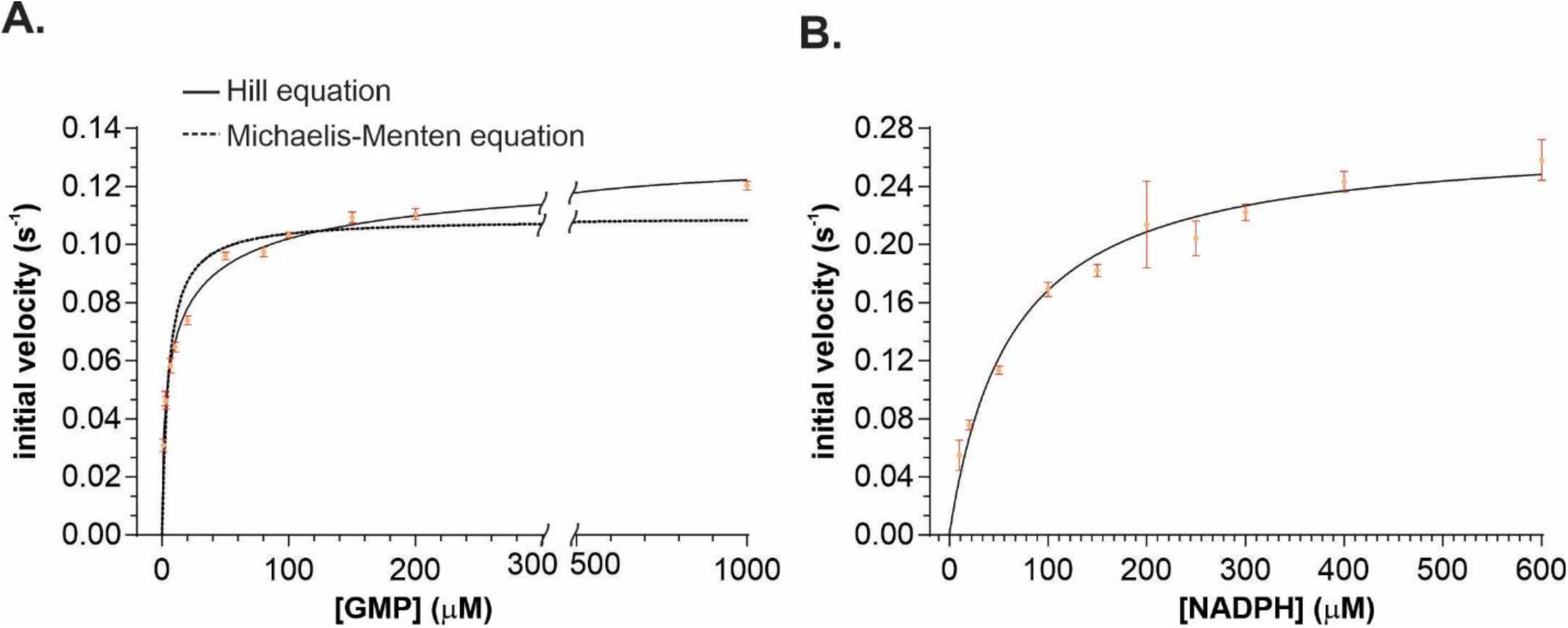
Steady state kinetics of the Mtb GMPR-catalyzed reaction at pH 7.6. Reaction rates were measured in 80 mM MPH buffer, 100 mM KCl and 20 nM Msm GMPR at pH 7.6 and 25 °C. **A)** Dependence of the initial reaction rate on the GMP concentration at fixed NADPH concentration (200 μM). Tha data were fitted with Hill (full line) or Michaelis-Menten (solid line) equations. The inner graph is a magnification of the grey area. **B)** Dependence of the initial reaction rate on the NADPH concentration at fixed GMP concentration (100 μM). The data were fitted with the Michaelis Menten equation.

**Fig S4.**
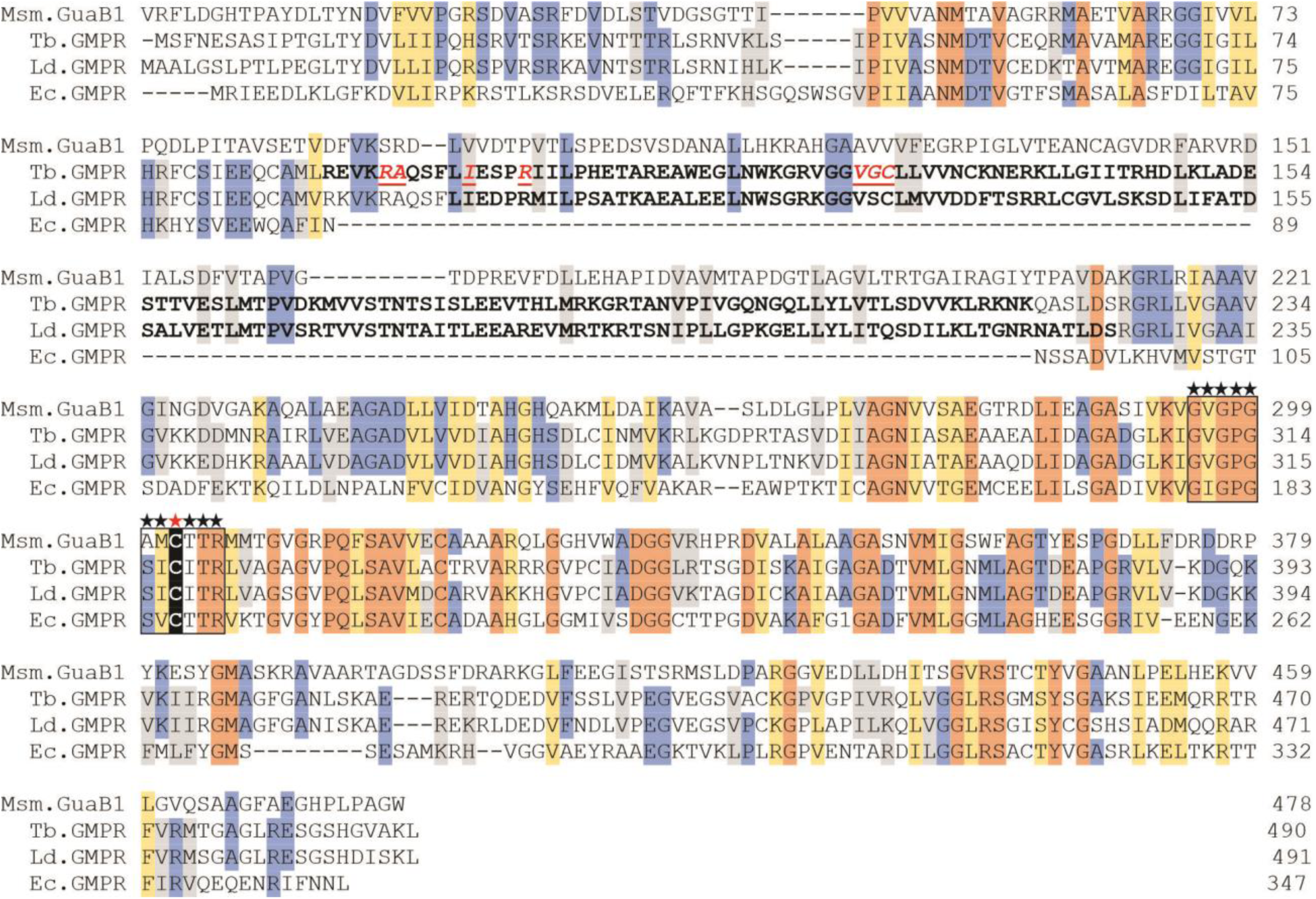
GuaB1 primary structure comparison with protozoal and *E. coli* GMPRs. The *Mycobacterium smegmatis* GMPR (RefSeq # WP_011729218.1) sequence was compared with those of GMPRs from *Trypanosoma brucei* (Tb.GMPR, RefSeq # XP_844882.1L), *Leishmania donovani* (Ld.GMPR, RefSeq # XP_001464746.1) and *Escherichia coli* (Ec.GMPR, RefSeq # WP_126739036.1) using Clustal X software. Identical and similar residues in all four proteins are shown in dark and light orange boxes, respectively. Identical and similar residues in three proteins are highlighted blue and gray, respectively. In Tb GMPR, the amino acid residues in bold letters represent the CBS domain, and the residues in red are involved in ATP binding (Imamura *et al*, 2020). In Ld GMPR, the amino acid residues in bold letters form the putative CBS domain proposed by (Smith *et al*, 2016). Amino acid residues of the catalytic core are framed and marked with stars. The catalytic cysteine residue is marked with a red star.

**Fig. S5.**
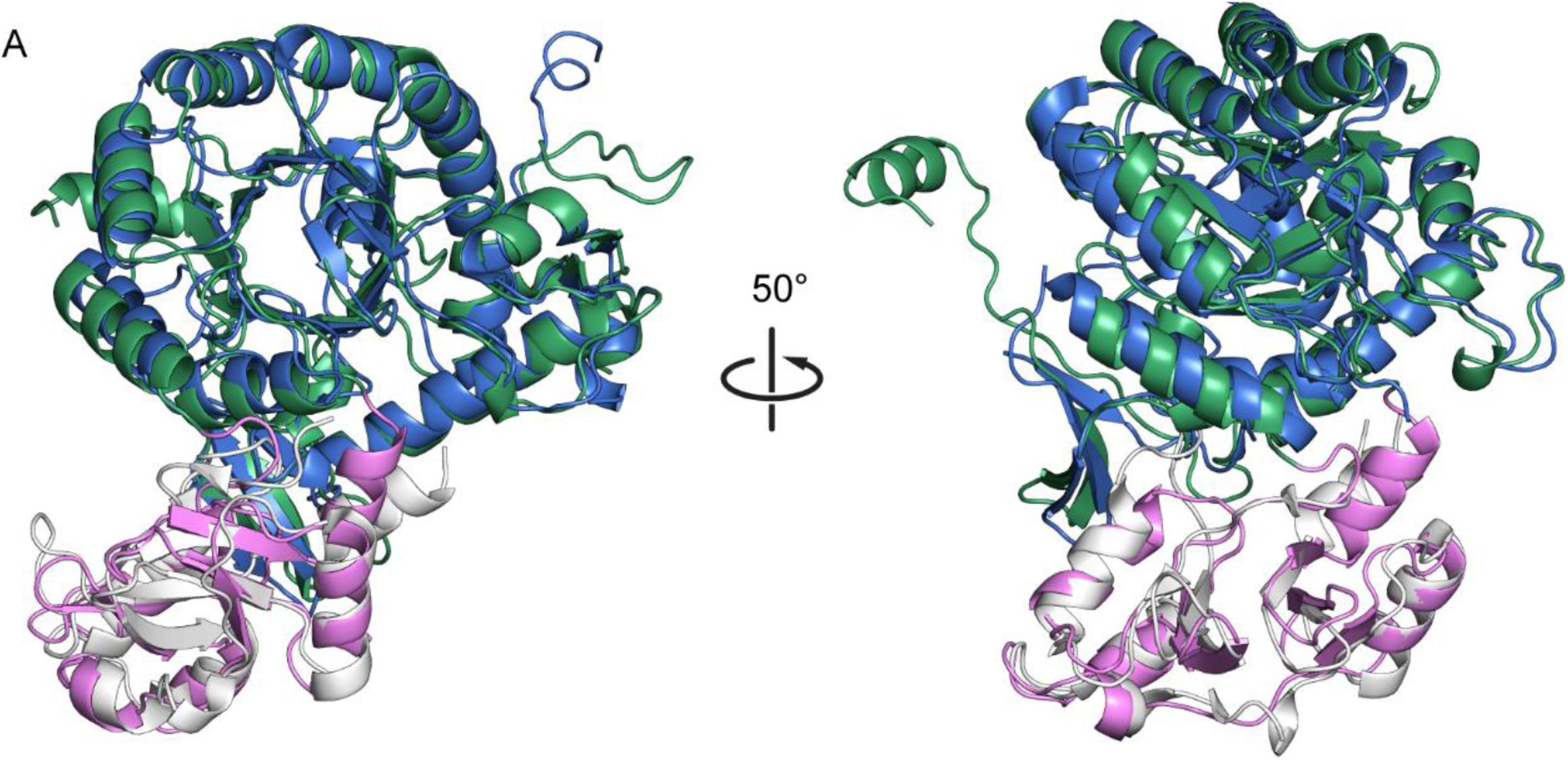
Comparison of Msm GMPR monomer (PDB ID 7OY9) and Tb GMPR monomer (PDB ID: 6JL8) structures. The catalytic (blue) and CBS (white) domains of Tb GMPR were aligned to Msm GMPR catalytic (green) and CBS (pink) domains independently, to compensate for differences in the quaternary structure.

**Fig S6.**
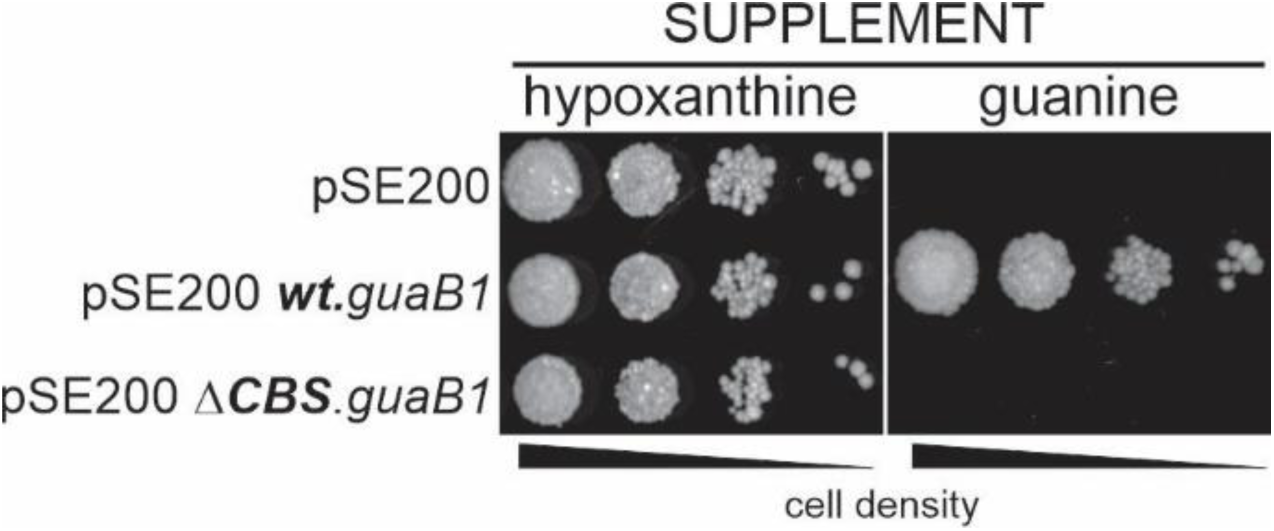
Importance of the CBS domain for Msm GMPR. The Δ*guaB1*Δ*purF* Msm strain was transformed with an empty pSE200 plasmid or pSE200 plasmid carrying genes for wt and Msm ΔCBS GMPR variants. Transformants were analyzed for growth in the absence and presence of 100 μM hypoxanthine or guanine on 7H10 medium containing 20 μg/ml kanamycin.

**Fig S7.**
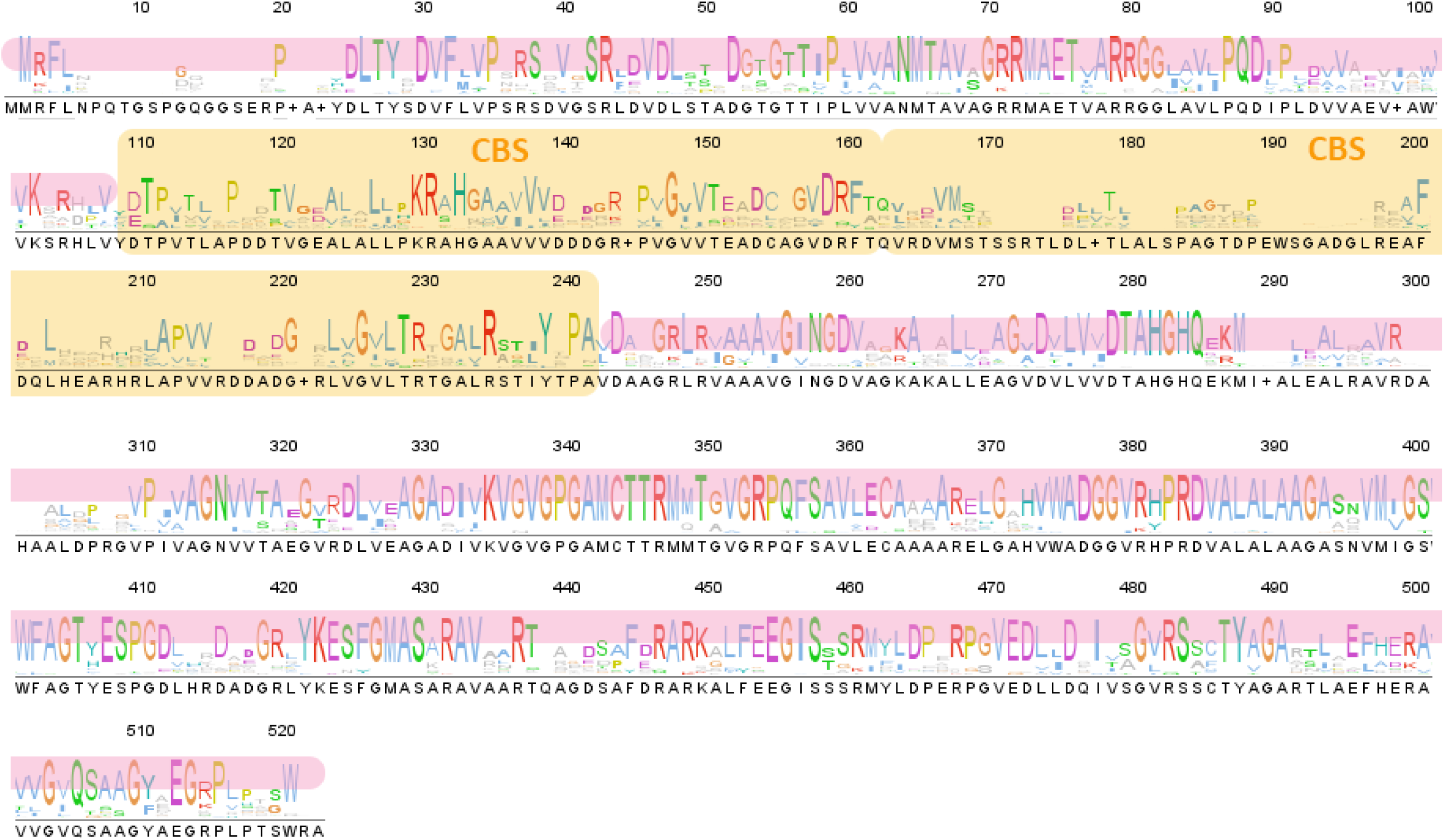
Conservation of GuaB1 across the *Actinobacteria* phylum. The height of the letters indicates the level of amino acid conservation; the colour of individual letters follows the ClustalX convention. The consensus of sequences aligned by the MUSCLE algorithm used for building a phylogenetic tree has the same domain composition as the *M. smegmatis* GuaB1 protein sequence. Pink and yellow shading indicate the catalytic and CBS domains, respectively.

**Supplementary table 1:**
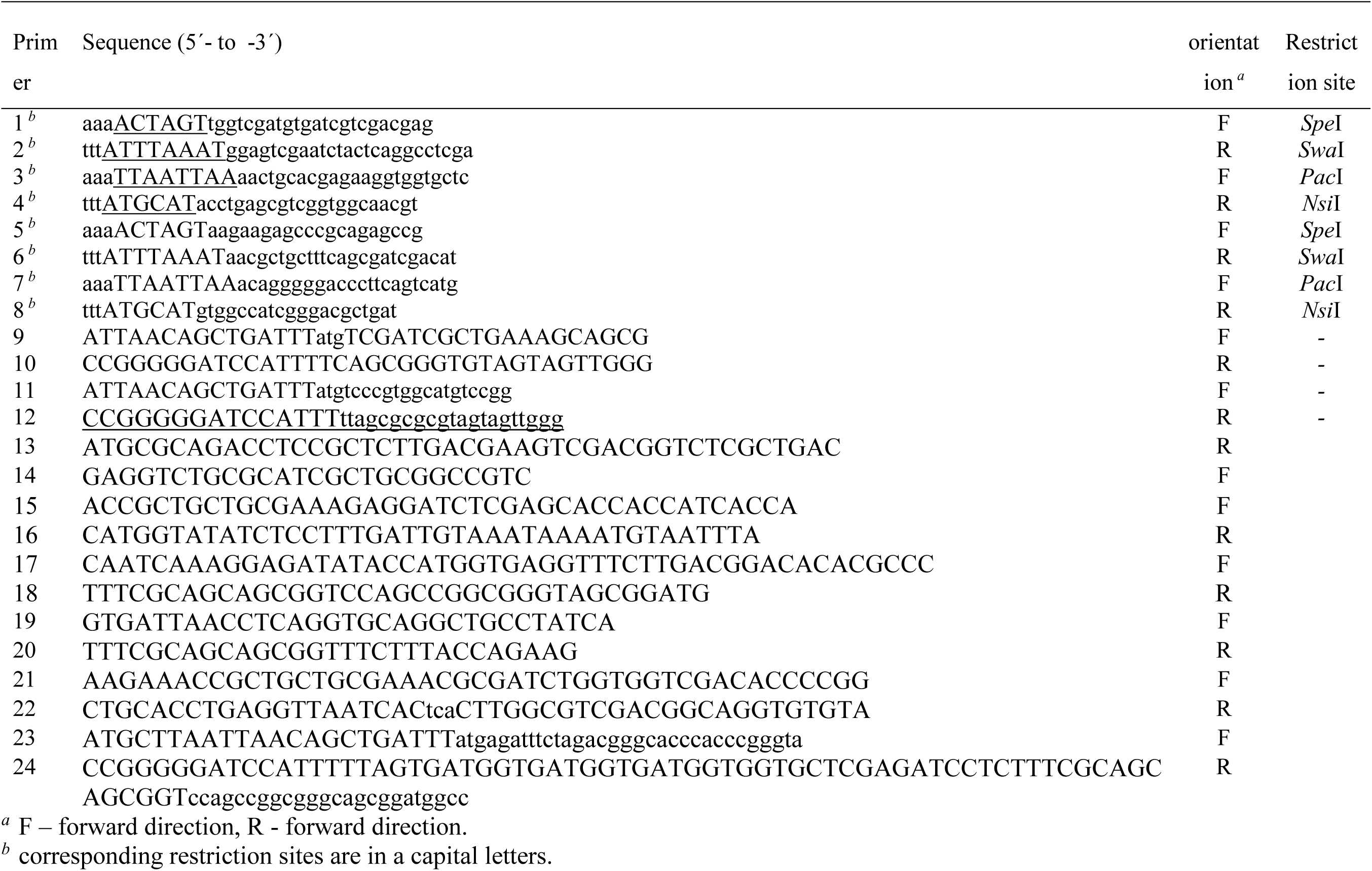
List of primers used in this study.

**Supplementary table 2.** Data collection and refinement statistics. Values for the highest resolution shell are shown in parenthesis

|  |  |
| --- | --- |
|  | Msm GMPR |
| PDB ID | 7OY9 |
| Data collection |  |
| Space group | C2 |
| Unit cell |  |
| a, b, c (Å) | 253.06, 109.83, 200.82 |
| $\alpha$, $\beta$, $\gamma$ (°) | 90.00, 119.43, 90.00 |
| Wavelength (Å) | 0.9184 |
| Resolution range (Å) | 43.73–2.8 (2.9–2.8) |
| Total reflections | 4685585 (479648) |
| Unique reflections | 118145 (11722) |
| Multiplicity | 39.7 (40.9) |
| Completeness (%) | 99.89 (99.94) |
| R-merge | 0.1555 (3.968) |
| R-meas | 0.1575 (4.018) |
| Average $I/\sigma(I)$ | 17.77 (1.07) |
| Wilson B-factor (Å<sup>2</sup>) | 94.66 |
| CC1/2 | 1 (0.662) |
| CC* | 1 (0.892) |
| Refinement |  |
| Resolution range (Å) | 43.73–2.8 (2.9–2.8) |
| Reflections used in refinement | 118056 (11718) |
| R-work | 0.2410 (0.4045) |
| R-free | 0.2791 (0.4468) |
| RMSD bond length (Å) | 0.010 |
| RMSD angle (°) | 1.25 |
| Atoms in AU | 25837 |
| Protein atoms in AU | 25837 |
| Protein residues in AU | 3647 |
| Average B-factor (Å<sup>2</sup>) | 114.34 |
| Ramachandran favored (%) | 98.42 |
| Ramachandran allowed (%) | 1.58 |
| Ramachandran outliers (%) | 0 |
| Rotamer outliers (%) | 1.43 |
| Clashscore | 5.28 |

